# A scalable, GMP-compatible, autologous organotypic cell therapy for Dystrophic Epidermolysis Bullosa

**DOI:** 10.1101/2023.02.28.529447

**Authors:** Gernot Neumayer, Jessica L. Torkelson, Shengdi Li, Kelly McCarthy, Hanson H. Zhen, Madhuri Vangipuram, Joanna Jackow, Avina Rami, Corey Hansen, Zongyou Guo, Sadhana Gaddam, Alberto Pappalardo, Lingjie Li, Amber Cramer, Kevin R. Roy, Thuylinh Michelle Nguyen, Koji Tanabe, Patrick S. McGrath, Anna Bruckner, Ganna Bilousova, Dennis Roop, Irene Bailey, Jean Y. Tang, Angela Christiano, Lars M. Steinmetz, Marius Wernig, Anthony E. Oro

## Abstract

**Background:** Gene editing in induced pluripotent stem (iPS) cells has been hailed to enable new cell therapies for various monogenetic diseases including dystrophic epidermolysis bullosa (DEB). However, manufacturing, efficacy and safety roadblocks have limited the development of genetically corrected, autologous iPS cell-based therapies.

**Methods:** We developed Dystrophic Epidermolysis Bullosa Cell Therapy (DEBCT), a new generation GMP-compatible (cGMP), reproducible, and scalable platform to produce autologous clinical-grade iPS cell-derived organotypic induced skin composite (iSC) grafts to treat incurable wounds of patients lacking type VII collagen (C7). DEBCT uses a combined high-efficiency reprogramming and CRISPR-based genetic correction single step to generate genome scar-free, *COL7A1* corrected clonal iPS cells from primary patient fibroblasts. Validated iPS cells are converted into epidermal, dermal and melanocyte progenitors with a novel 2D organoid differentiation protocol, followed by CD49f enrichment and expansion to minimize maturation heterogeneity. iSC product characterization by single cell transcriptomics was followed by mouse xenografting for disease correcting activity at 1 month and toxicology analysis at 1-6 months. Culture-acquired mutations, potential CRISPR-off targets, and cancer-driver variants were evaluated by targeted and whole genome sequencing.

**Findings:** iPS cell-derived iSC grafts were reproducibly generated from four recessive DEB patients with different pathogenic mutations. Organotypic iSC grafts onto immune-compromised mice developed into stable stratified skin with functional C7 restoration. Single cell transcriptomic characterization of iSCs revealed prominent holoclone stem cell signatures in keratinocytes and the recently described Gibbin-dependent signature in dermal fibroblasts. The latter correlated with enhanced graftability. Multiple orthogonal sequencing and subsequent computational approaches identified random and non-oncogenic mutations introduced by the manufacturing process. Toxicology revealed no detectable tumors after 3-6 months in DEBCT- treated mice.

**Interpretation:** DEBCT successfully overcomes previous roadblocks and represents a robust, scalable, and safe cGMP manufacturing platform for production of a CRISPR-corrected autologous organotypic skin graft to heal DEB patient wounds.

## INTRODUCTION

Over the past decade, advances in the field of stem cell biology and regenerative medicine have enabled the prospect of genetically corrected autologous tissue replacement for previously untreatable conditions. Past clinical successes have mainly used viral gene transfer into somatic tissue, such as the bone marrow stem cells, illustrating that genetic correction of stem cells capable of tissue regeneration provides long-term disease modifying activity ^1, 2^. Somatic cell reprogramming into induced pluripotent stem (iPS) cells allows the generation of patient-derived, and thus autologous, iPS cells that can be genetically manipulated. iPS cells can be differentiated into not only individual cell types, but also organotypic cultures or organoids containing multiple key cell types that compose a tissue. However, such complex, multi-lineage manufacturing methods have not yet been developed at scale for clinical evaluation ^3–5^. The iPS cell-based approach provides a solution for the two main limitations of current somatic cell and gene therapy strategies: (i) iPS cells can be grown to virtually unlimited numbers providing a solid foundation for tissue organoid production-scalability and (ii) defined gene editing recreates a wild type allele while avoiding retroviral insertional mutagenesis. Thus, the combination of cell reprogramming, genomic correction of pathogenic mutations and composite cell transplantation has the potential to eradicate the impacts of disease-causing mutations in afflicted tissues ^5–7^.

While appealing in early studies, translation of corrected autologous iPS cell-derived products from proof-of-concept towards realistic clinical manufacturing has been met with a panoply of technical and regulatory roadblocks. Hurdles include developing a robust and reproducible manufacturing method that overcomes critical bottlenecks, including iPS cell generation, validated genetic correction, and safe and effective differentiation into desired tissues. A second set of hurdles includes sequential cell manipulation that results in protracted and labor-intensive manufacturing, increasing batch variability and compromising genomic integrity ^8^. Moreover, open questions regarding regulatory concerns, including determination of the safety risk of a pluripotent cell-derived product, avoidance of animal-derived products, and a risk assessment of genetic mutations introduced during cell culture and genetic engineering, have hampered wide- spread adoption of this cell/tissue therapeutic platform ^9, 10^.

The blistering disorder Dystrophic Epidermolysis Bullosa (DEB) maps to mutations in the *COL7A1* gene and results in extreme skin fragility due to collagen VII loss at the basement membrane zone (BMZ) ^11–13^. Without curative treatment, the only option remains palliative wound care. Painful chronic wounds severely impact quality of life and the chronic inflammatory milieu of constantly re- and degenerating skin wounds invariably results in the formation of an aggressive form of squamous cell carcinoma to which most patients ultimately succumb ^14–16^. Disease severity and the lack of treatment options motivates the development of scalable and safe manufacturing options for autologous tissue replacement technologies ^17–19^. Increasing efforts towards cell and gene therapies to treat Junctional EB have been led to great promise with the basal layer of the skin being self-renewing and demonstration that correcting the basal keratinocytes that contain holoclones with long-term stem cell activity has remarkable disease- modifying potential ^17, 20, 21^. In addition, a recent Phase I/IIA trial demonstrated that autologous grafts of expanded somatic RDEB keratinocytes transduced with a retroviral delivered *COL7A1* cDNA has highly efficient wound healing capability ^22^. While these groundbreaking clinical trials showed disease-modifying activity, the approaches have important limitations including difficulty to reliably expand somatic RDEB keratinocytes and the safety concern of insertional mutagenesis by retroviral gene transfer ^23, 24^. Development of clinically scalable iPS cell-derived skin replacement that overcomes current manufacturing challenges would represent a major advance for many genetic diseases, including DEB.

Here, we realize the advantages of an iPS cell-based multi-lineage differentiation approach to generate organotypic composite grafts via next generation genetic and cellular engineering. Solving critical bottlenecks, we refine a practical and simplified Good Manufacturing Practice (GMP)-compatible protocol for the generation of genetically corrected autologous organotypic skin grafts that include keratinocytes, dermal fibroblasts, and melanocytes for the long-term healing of DEB patient wounds.

## RESULTS

### Optimization of CRISPR/CAS9-mediated targeting of the *COL7A1* locus

While previous work ^8, 25, 26^ demonstrated the possibilities of *ex vivo* autologous iPS cell-based gene therapy for treatment of Recessive Dystrophic Epidermolysis Bullosa (RDEB), several hurdles preventing clinical translation remained. These include: 1) relatively inefficient iPS cell derivation and genetic correction, and 2) lack of defined and efficient protocols for differentiation of edited iPS cells into multi-lineage induced skin composites (iSCs). Consequently, the previous protocols took many months to complete and involved multiple clonal steps, greatly increasing complexity and procedural variabilities, thereby complicating the development of Standard Operating Procedures (SOPs) while pronouncing the rate of culture-induced mutations^8^. To overcome these limitations, we evolved a next-generation, scalable, non-integrating, xeno-free, and GMP-compatible platform that produces an epidermal-dermal-melanocyte containing, organotypic iSC product for long-term patient wound healing.

We first designed a cGMP method that allows derivation of *COL7A1*-corrected iPS cells from primary patient fibroblasts in 4 weeks. This SOP integrates iPS cell reprogramming and gene correction into a single manufacturing step (Fig. 1A), reducing culture time, mutational burden and clonal bottlenecks. In this design, primary patient fibroblasts from a dermal punch biopsy are transiently transfected with (i) CAS9-sgRNA containing ribonucleoproteins (RNPs), (ii) single stranded oligodeoxynucleotides (ssODNs) that encode the desired genomic correction, and (iii) reprogramming factors-encoding mRNAs that induce iPS cells. iPS cell colonies, emerging ∼14 days after the initial transfection with reprogramming factors, are then isolated and screened via droplet digital (dd) polymerase chain reaction (PCR) employing probes specific for the properly corrected *COL7A1* locus. Ensuing quality controls validate *COL7A1* edits, cellular identity, and genomic/chromosomal stability.

**Figure 1:**
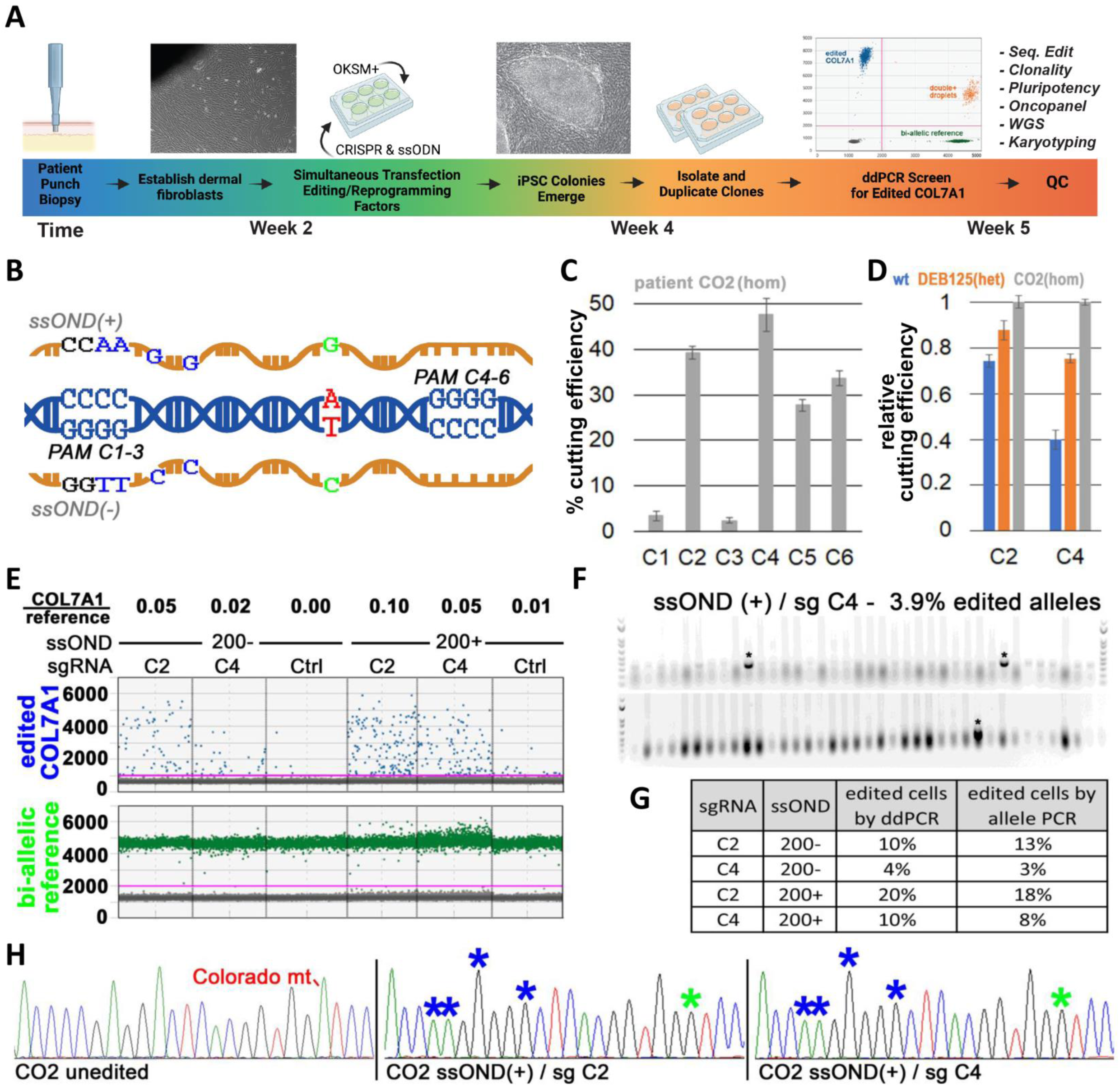
Optimized editing of the *COL7A1* Colorado mutation (7485+5 G>A). **(A)** Overview of single editing&reprogramming manufacturing step. **(B)** Overview of the Colorado mutation (red) and (+) or (-) strand ssODNs used for gene-editing. 6 possible PAM sites (C1-6) can be engaged via sgRNAs to mediate cutting of pathogenic alleles via CRISPR/CAS9. ssODNs encode for 4 silent mutations (blue) and the wild type sequence (green). **(C)** Absolute CRISPR/CAS9 cutting efficiencies of the homozygous Colorado mt in CO2 patient fibroblasts as mediated by sgRNA C1-C6 (n=2; stdev). **(D)** Relative CRISPR/CAS9 cutting efficiencies of the homozygous (CO2), heterozygous (DEB125) Colorado mt or the wild type locus as mediated by sgRNA C2 or C4 in indicated patient fibroblasts (n=2; stdev). **(E)** *COL7A1* editing efficiencies measured by ddPCR in CO2 primary patient fibroblasts after transfection with ssODNs and sgRNA/CAS9-containing RNPs as indicated. A bi-allelic locus (green) is used as a reference for calculating *COL7A1* editing (blue) efficiencies. Both, strandedness of the ssODN and sgRNA affect editing efficiencies. Ctrls omitted sgRNAs. **(F)** Agarose gels visualizing 77 E. coli colony- PCRs to detect edited Topo-cloned *COL7A1* alleles from cells treated as in (E) with ssODN/sgRNA combinations as indicated (see Fig. S1 for remaining ssODN/sgRNA combinations). A primer specific for intended silent mutations (see Fig. 1B) only yields PCR products of alleles with integration of donor sequences (asterisks). **(G)** Summary of *COL7A1* editing efficiencies achieved with different ssODN/sgRNA combinations as measured by ddPCR (Fig. 1E) and as observed by Topo-cloning of individual alleles (Fig. 1F). Mono-allelic integrations of ssODN sequences were assumed for calculations (see discussion). **(H)** Sanger sequence traces of unedited and edited alleles from (Fig. 1F, S1H). Asterisks indicate integration of intended silent mutations (blue) and repair of the pathogenic mutation (green).

To test the applicability of this approach, we assessed the potential for gene targeting of the *COL7A1* locus in patient-derived dermal fibroblasts from three individuals carrying the so-called “Colorado” mutation, i.e. *COL7A1* c.7485+5G>A (Fig. 1B). Patients CO1 and CO2 carry homozygous and patient DEB125 carries a compound heterozygous “Colorado” mutation (*COL7A1* c.6527dupC is the other pathogenicity of DEB125). Initially we tested all possible sgRNAs mediating CAS9-cutting of the Colorado-allele and ssODNs of various lengths encoding for either the (+) or (-) strand of DNA (Fig. 1B). While the 6 possible sgRNAs specific to the Colorado mutation exhibited favorable *in silico* predicted specificity and activity scores (Fig. S1A), Tracking of Indels by Decomposition (TIDE) analysis in patient fibroblasts transfected with RNPs showed variable efficiencies (Figs. 1C, S1B-C). Focusing on the two most efficient sgRNAs (i.e. sgRNA C2 and C4), we analyzed their specificity for the Colorado allele (Fig. 1D). TIDE analysis of wild type (wt), heterozygous DEB125, and homozygous CO2 fibroblasts transfected with RNPs revealed that sgRNA C4, with a protospacer adjacent motif (PAM) closer to the Colorado mutation, is more specific for the disease allele.

Next, we optimized sgRNA C2 and C4 RNP-mediated repair of mutant *COL7A1* using ssODNs encoding for the wt *COL7A1* sequence and 4 silent mutations used for detection of editing events via specific ddPCR probes (Fig. 1B). By comparing with a bi-allelic reference locus, ddPCR allowed quantification of the edited *COL7A1* allele. This approach showed that sgRNA C2 mediated 2-2.5x more *COL7A1* targeting events than sgRNA C4 (Fig. 1E). The ssODN length (up to 200 nt) positively correlated with editing efficiencies (Fig. S5B-C). Surprisingly, also the strandedness of ssODNs influenced editing efficiencies with the (+) strand-encoding sequence yielding 2-2.5x higher efficiencies than the (-) ssODN (Fig. 1E) in both homozygous and heterozygous patients (Fig. S1D). We confirmed the ddPCR results by cloning the target locus of bulk-edited fibroblasts into plasmids, followed by analysis of individual, cloned alleles via PCR primers specific for the edited locus (Figs. 1F, S1F-H). Analysis of 77 individual target alleles each, from cells treated with all possible combinations of sgRNA C2 or C4 and ssODN (+) or (-) validated the ddPCR results. Sanger sequencing revealed that both, sgRNA C2 and sgRNA C4 are mediating *COL7A1* editing as intended when used in combination with (+) ssODNs (Figs. 1G-H, S1F-H). Remarkably, although all the Sanger-sequenced alleles from cells treated with (-) ssODNs exhibited at least partial integration of donor sequences, none of them was correctly edited (not shown). We conclude that all parameters involved, i.e. the sgRNA sequence, the length and strandedness of ssODNs, and the particular target locus must be tested for optimal efficiency and specificity of editing events.

### Combining iPS cell-reprogramming and *COL7A1* correction in one manufacturing step

The optimized *COL7A1* targeting efficiency allowed us to test whether it may be feasible to correct and reprogram cells in a single manufacturing step, which would greatly minimize production time and eliminate multiple clonal selections. We chose to deliver the reprogramming factors via transfection of mRNA since this approach has been shown to be efficient, is compatible with GMP manufacturing using chemically defined reagents, and represents a transient treatment leaving no genetic scars ^26^. In line with the TIDE assay in fibroblasts (Fig.1D), sgRNA C2-mediated *COL7A1*-editing in DEB125 fibroblasts directly followed by mRNA reprogramming resulted in ∼15% targeted iPS cell colonies but was not specific for repairing the mutant allele of this heterozygous patient (Fig. S2). We therefore focused on sgRNA C4, which mediates only slightly lower editing efficiency but is more specific for the Colorado allele. First, we determined via a dose range that 5 pmole sgRNA C4-containing RNP per 30k fibroblasts was sufficient for optimal cutting of the Colorado allele (Fig. S1E). Next, we measured the *COL7A1* editing efficiencies in fibroblasts of patients CO1 and CO2, which carry a homozygous Colorado mutation, using engineered high fidelity CAS9s. Reassuringly, ddPCR, using ssODN (+) and sgRNA C4 in complex with CAS9s HiFi or SpiFy showed targeting events in 3-7% of the cells (Fig. 2A-B). These encouraging efficiencies prompted us to test induction of reprogramming immediately following *COL7A1* editing via GMP-compatible SpiFy CAS9. We screened 186, 293, and 24 iPS cell clones derived from patient CO1, CO2 and DEB125 by ddPCR, yielding 8, 25, and 1 candidate lines, respectively (Fig. 2C, G and S4). Analysis of 5, 11, and 1 candidate iPS cell lines from each patient via conventional PCR amplification of the target locus, followed by plasmid cloning and Sanger sequencing of individual alleles, revealed that all but 2 lines were correctly edited (Fig. 2D-G). As expected from the specificity and efficiency mediated by guide C4 (Fig. 1D), in iPS cell lines from patients homozygous for the Colorado mutation (i.e. CO1 and CO2) the second allele acquired InDel mutations of various sizes in all cases, whereas the second, wt allele in iPS cells derived from the heterozygous patient DEB125 remained unperturbed. Of important note, in some iPS cell clones derived from homozygous Colorado patients the InDels on the second allele were larger than the 731bp PCR amplicon that we initially chose for analysis (Fig. 2D), wrongly implying bi-allelic *COL7A1*- correction. Larger PCR amplicons identified some of these bigger InDels, e.g. a 654bp deletion in line CO2-65(B), while the nature of others (e.g. line CO1-48) could not be determined (Fig. 2E-G; see discussion).

**Figure 2:**
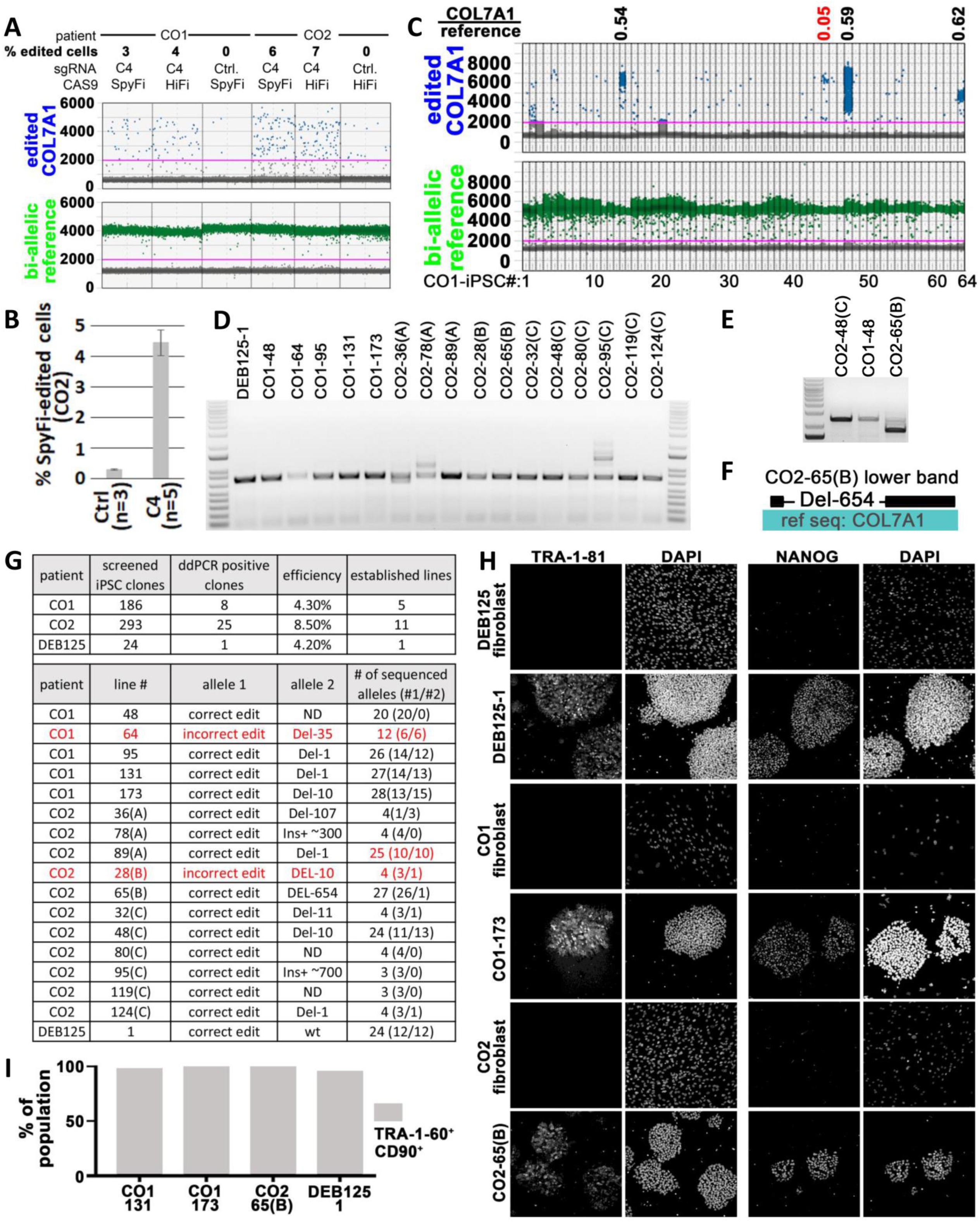
Successful single manufacturing step editing/reprogramming of patients CO1, CO2 & DEB125. **(A)** *COL7A1* editing efficiencies as measured by ddPCR in CO1 and CO2 primary patient fibroblasts (homozygous Colorado mt) after transfection with (+)ssODN and RNPs containing sgRNA C4 and high fidelity CAS9 HiFi or SpyFi as indicated. A bi-allelic locus (green) is used as a reference for calculating *COL7A1* editing (blue) efficiencies, assuming mono-allelic editing events. Ctrls omitted sgRNAs. **(B)** Reproducible *COL7A1* editing in CO2 primary patient fibroblasts as in (Fig. 2A) with SpyFi CAS9 (error bars are sem). **(C)** Representative ddPCR screen of 64 single-step edited/reprogrammed iPS cell lines derived from patient CO1 fibroblasts. Ratios of signals detecting edited *COL7A1* alleles (blue) and a bi- allelic reference locus (green) are used to identify mono- (0.5 +/-0.19) or bi-allelic (1.0 +/-0.19) editing events (black values; red values below/above cut off indicate potentially mixed or incorrectly edited clones). **(D-E)** Agarose gel visualizing PCR amplicons of a 731bp (D) and 2418bp (E) sequence surrounding the edited *COL7A1* locus from single-step edited/reprogrammed iPS cell lines derived from 3 patients heterozygous (DEB125) and homozygous (CO1 and CO2) for the Colorado mt. Note that some samples yield 2 PCR products, indicative of large InDels on one of the *COL7A1* alleles. InDels can be substantial (e.g. line CO2-65(B)), so that they are only included on bigger PCR products (E). See text for details. **(F)** Sanger sequencing of the smaller PCR product from line CO2-65(B) from (Fig. 2E) reveals a large 654bp deletion when aligned with the wt COL7A1 sequence. This deletion is not detected by smaller PCR amplicons in (Fig. 2D). **(G)** Summary of single-step editing/reprogramming screens conducted with sgRNA C4 and (+)ssODN from 3 patients as indicated (top). Topo-cloning and sanger sequencing of PCR products from (Fig. 2D-F) confirms correct COL7A1 editing on target alleles in 15 of 17 single-step edited/reprogrammed iPS cell lines. **(H)** Immuno-fluorescence microscopy images of single-step edited/reprogrammed iPS cells and parental fibroblasts stained for pluripotency markers TRA-1-81 and NANOG from 3 patients. DAPI visualized DNA. **(I)** Summary of flow cytometry analysis of single-step edited/reprogrammed iPS cells from 3 patients for CD90 and the pluripotency marker TRA-1-60 (see Fig. S3D for details).

Next, we chose candidate iPS cell lines for deeper genomic characterization. Sanger sequencing of up to 28 individual target loci cloned into plasmids showed an equal distribution between correctly edited and the second allele in all but one sample (i.e. CO2-89(A); Fig. 2G). In iPS cell lines derived from homozygous patients, the second allele that did not incorporate the ssODN always displayed a characteristic InDel, indicating high efficiency of employed RNPs. Other than iPS cell line CO2-89(A), in which 20% of sequences displayed deletions of a different nature (not shown), our data is consistent with clonal origin of picked iPS cell lines. In addition, we verified cellular identity of our iPS cell lines by robust expression of the pluripotency markers TRA-1-81, TRA-1-60, NANOG, *Lin28*, *Oct4,* and *Sox2* as determined by immunofluorescence, cytometry, and/or RT-PCR (Fig. 2H-I, S3D, S6A).

Finally, to test applicability to correct other mutations, we followed the same SOPs developed for the Colorado mutation and successfully derived one-step corrected iPS cell lines with similar efficiency from a fourth patient, DEB135, who carries two different compound heterozygous mutations, i.e. *COL7A1* c.6781C>T and c.6262 G>A (Fig. S5). Importantly, this demonstrates that our single clonal step iPS cell manufacturing process can in theory be adapted for other therapeutic cell manufacturing procedures.

### Scalable and reproducible differentiation of DEB iPS cells into organotypic skin grafts

Proper mammalian skin development requires the interaction between early surface ectoderm progenitors, regional mesoderm, and neuroectoderm ^27–30^. We recently demonstrated the importance of a subtype of mesoderm that depends upon the chromatin regulator Gibbin for proper epidermal stratification ^31^. We therefore sought to develop a novel multi-lineage cutaneous organoid differentiation method that imitates the interaction and co-dependence of the cell lineages that cooperate in the embryo to form skin (Fig. 3A) ^31, 32^. First, embryonic surface ectoderm was induced with retinoic acid/bone morphogenic protein-4 (RA/BMP4) for 7 days ^31^. scRNA-seq analysis of day 7 cultures showed the successful creation of surface ectoderm, mesoderm, and neuroectoderm (Fig. 3B,C). The second inductive phase used a defined matrix and media containing the epidermal growth factors insulin, EGF and FGF for an additional 40-45 days, allowing reciprocal epithelial-mesenchymal-neuroectodermal interactions to mature the cultures into a therapeutic organotypic induced skin composite (iSC).

**Figure 3:**
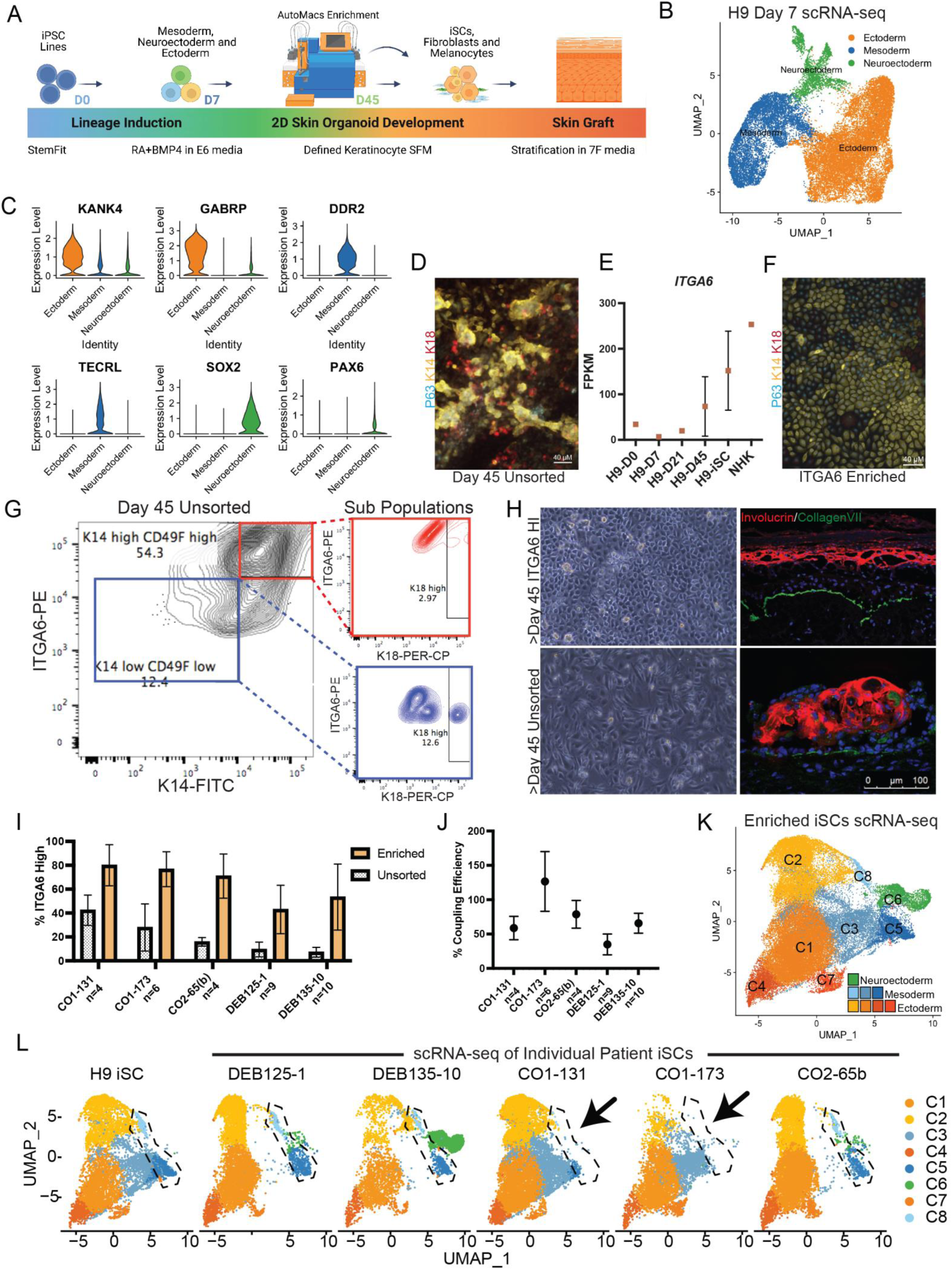
Generation of organotypic skin grafts at clinical scale. **(A)** Differentiation strategy and cell enrichment with AutoMACS pro-separator for iPS cell clone comparison following a defined GMP compatible protocol to manufacture induced skin composite (iSC) grafts. **(B)** UMAP cluster representation of scRNA sequencing of H9 ES cell differentiation at Day 7 reveals the three lineage clusters: ectoderm (orange), mesoderm (blue) and neuroectoderm (green). **(C)** Violin plots depicting RPKM relative expression of representative ectoderm, mesoderm and neuroectoderm genes. **(D)** Immuno-fluorescence (IF) of iPS cell line DEB135-10-derived iSCs, stained for p63, K18 and K14 at day 45 of differentiation as in (Fig. 3A). **(E)** ITGA6 expression as per RNA-seq in H9 ES cells during iSC differentiation. Time points are indicated (D: day). NHKs were used as a positive control. **(F)** IF staining of AutoMACS-ITGA6 enriched, expanded iSCs. p63 (Cyan), K18 (red), and K14 (yellow). **(G)** Flow cytometry of day 45 unsorted H9 ES cells. Cells highly double positive for ITGA6 PE (y-axis) and K14 FITC (x-axis) are in high gate (red) and lower ITGA6/K14 expressing cells are in low gate (blue) to evaluate expression of K18 (PER-CP) in the subpopulations (right). **(H)** (Top left) Bright field image of FACS sorted ITGA6^HI^ H9 ES cell-derived iSCs expanded to day 50 and used for organotypic stratification (Top right). IF shows normal polarization and stratification of the epidermis from the ITGA6^HI^ H9 ES cell-derived iSCs (Collagen 7 (green), Involucrin (red), DAPI (blue)); (Bottom left) bright field image of unsorted H9 ES cell derived iSCs used for organotypic stratification (Bottom right). IF of corresponding unsorted H9 cells revealed disorganized layering and stratification. **(I)** Flow cytometry analysis of %ITGA6 positivity measured before and after AutoMACS enrichment for 5 differentiated iPS cell patient lines (independent differentiation runs for each clone n=4-10). **(J)** % Coupling efficiency (CE) determined by the equation %CE= live sorted iSCs/iPS cell input*100. **(K)** UMAP showing overlay of scRNAseq datasets from (Fig. 3L) with color scheme as indicated. 4 ectoderm (C1, C2, C4, C7), 3 mesoderm (C3, C8, C5) and 1 melanocyte / neuroectoderm-derived (C6) clusters were identified. **(L)** UMAP of scRNA sequencing of 5 individual patient iSCs and H9 ES cell derived iSC after ITGA6 enrichment and expansion for 2 passages *in vitro*, revealing 8 clusters (C) comprising the DEBCT product. Gibbin dependent clusters are outlined in dotted lines (see text for details), and arrows show lack of these clusters in products derived from iPS cell lines C01-131/173.

A common hurdle in pluripotent cell differentiation comes from stochastic mechanisms during complex cell culture that lead to variable keratinocyte maturation and presence of immature K14^+^, K18^+^ epithelial cells (Fig.3D) ^33^. To overcome this hurdle and to enrich for mature basal keratinocytes, we used the previously characterized stem cell surface marker ITGA6 /ITGB4 ^34, 35^. We verified that ITGA6 is expressed highly on p63^+^; K14^+^; K18^-^ cells, while K18^+^ cells were ITGA6 low or negative. An ITGA6 magnetic bead-based, automated pro-separator AutoMACS® efficiently enriched for p63^+^; K14^+^ cells and removed K18+ cells (Fig. 3E-G, and S6B-C). In contrast to unenriched populations, ITGA6-enriched cells produced robust stratified epidermis/dermis in liquid/air interface organotypic cultures as demonstrated by involucrin expression and deposition of collagen VII to the BMZ (Fig. 3G-H). Importantly, mesodermal and melanocytic cell populations were still present after ITGA6-enrichment allowing continued signaling between cell types (Fig. 3L,K). We successfully differentiated and AutoMACS®- enriched five independent, genetically corrected DEB patient cells lines in multiple replicates demonstrating reproducibility among different lines from different patients. The “coupling efficiency”, i.e. the ratio between ITGA6-enriched cells after differentiation and the input iPS cells, ranged between 40-130% (Fig. 3I, J), demonstrating the robustness of the process.

We next sought to implement ITGA6 enrichment at a clinical manufacturing scale. We performed 5 large scale differentiation runs that improved iSC formation by employing a CliniMACS® Plus cell separator in 3 different modes, yielding various enrichment and cell viability ratios. Limited cell expansion *in vitro* generated the needed cell numbers for future clinical trials (Fig. S6D-H). Flow cytometry and bulk RNA-seq verified successful enrichment (Fig. S6B,E,H), demonstrating the feasibility of clinical scale manufacturing.

Lastly, we used single cell transcriptomics to characterize the final post-enrichment iSC, which would be equivalent to the DEBCT clinical product (Fig. 3K,L). We profiled iSC cultures from H9 cells and 5 patient iPS cell lines, revealing 8 distinct cell clusters representing surface ectodermal (C1, C2, C4, C7), mesodermal (C3, C5, C8) and melanocyte cell types (C6) (Fig. S6I) ^31, 36–40^. C1 most closely resembles basal keratinocytes (high K14, low K18, low cell cycle markers), C2 resembles long-term proliferative “holoclone” keratinocyte stem cells (high CDK1 and TOP2A; Fig. S6I) ^41, 42^. Cells in C4 and C7 initiated signatures of the early epidermal stratification phase. C5 and C8 cells closely resemble Thy1/CD90+ Gibbin-dependent mature dermal fibroblasts that are required for epidermal induction ^31^, and the C3 cluster contains cells of more immature dermal / pre-vascular characteristics ^36^. Intriguingly, the 6 cell lines produced different ratios of these induced cell clusters, allowing us to correlate the functional implications of their presence (Fig. 5C-F).

### Genomic stability of *COL7A1-*corrected iPS cells and iSCs

A critical safety aspect of cell expansion is genomic and chromosomal stability, as we estimate that 3×10^7^ undifferentiated iPS cells are needed to generate one clinical iSC application of a 6x8 cm sheet graft in a Phase I/IIa trial (Fig.S6G ^19^). Unlike previously tested media ^8^, a more recently developed chemically defined media on plates coated with the E8 fragment of Laminin- 511 ^43^ allowed expansion of karyotypically normal iPS cell lines derived from 5 individuals with 3 sgRNAs to at least 3×10^7^ cells in 11 of 12 instances (Figs. 4A, S7A). As part of our product safety methodology, we performed whole genome sequencing (WGS, 40x coverage) of four single-step corrected iPS cell lines from three patients carrying the Colorado mutation, their parental fibroblasts and differentiated ITGA6-enriched iSCs. WGS data confirmed all CAS9- mediated *COL7A1* edits (Fig. 2G). To identify novel single nucleotide variants (SNVs) and insertions or deletions (InDels) with high confidence, we used the agreement between three variant callers and filtered out all SNVs/InDels previously annotated in SNV/InDel databases (see material and methods). This strategy yielded an average of 70,386 +/-570 (SEM) novel SNVs and 32,465 +/-1,506 (SEM) novel InDels in each of the 11 samples. To find SNVs and InDels that are present specifically in either iPS cells and/or iSCs and thus, may be induced and/or selected for by our manufacturing process, we employed two alternative and complementary strategies: (i) k-means clustering by allele frequency (AF) and (ii) AF/odds ratio cut-off filtering. The k-means clustering at high resolution (feature space k=9) showed that most SNVs/InDels exhibit similar AFs among all three cell types, suggesting that these variants are heterozygous or homozygous pre-existing germline variants (Fig. 4B and Fig. S7B). Only one cluster (cluster 7 for DEB125-1, Fig. 4B) contained iPS cell/iSC-specific variants and no cluster was identified with variants unique to either iPS cells or iSCs. The same pattern was observed in all four experiments. Importantly, there were only three iPS cell/iSC-specific variants found in more than one patient (Fig. S7C). Of these three variants, one shared between DEB125 and CO2 maps to an intron of *Magi2* and is already present in the respective parental fibroblasts at AF=0.1-0.2. Similarly, the two variants shared between lines CO2-65(B) and CO1-173, mapping to an intron of a ncRNA (i.e. LOC100507053) and an intergenic region, are also found in parental fibroblasts at AF=0.2-0.3, and AF=0.1-0.2, respectively. Thus, all three variants, themselves of unknown significance, were in fact already pre-existing in parental fibroblasts at lower AFs and are not introduced *de novo* by our manufacturing process.

**Figure 4:**
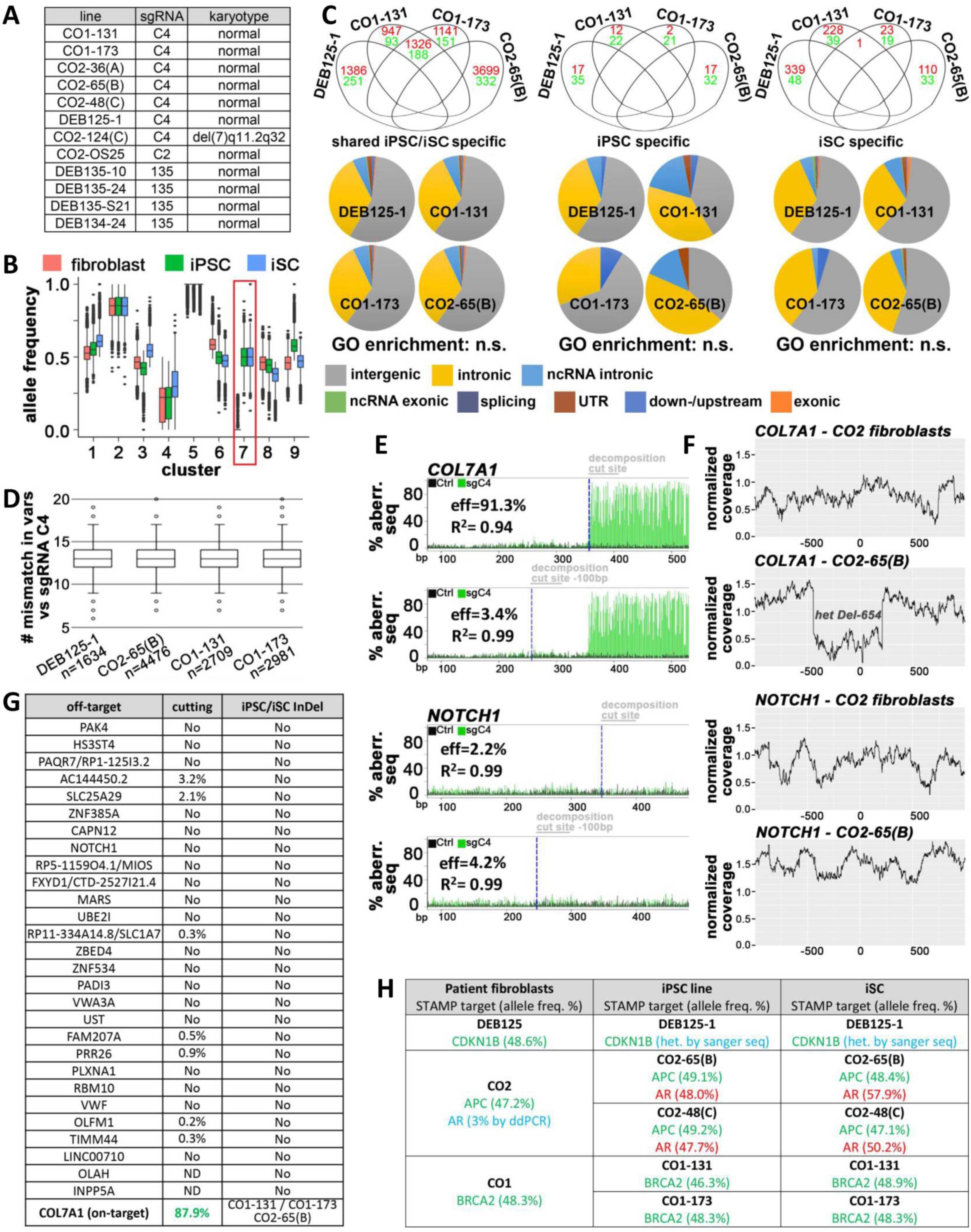
Genomic and chromosomal stability of single manufacturing step edited/reprogrammed iPS cells and iSCs. **(A)** Normal karyotypes were observed in 11 of 12 single-step edited/reprogrammed iPS cell lines from 5 patients (CO1, CO2, DEB125, DEB134, and DEB135). **(B)** K-means clustering of all novel variants (n=111741) found by 40x whole genome sequencing (WGS) in fibroblasts and thereof derived iPS cells and iSCs from patient DEB125 (see Fig. S7 for other patients). Only a small sub-set of variants (red frame) is found to have differential allele frequencies (AFs) in iPS cells/iSCs compared to parental fibroblasts. **(C)** A defined cut-off (AF=0.25), combined with odds ratio filtering (see material and methods) identifies variants from 40x WGS (red: SNPs; green: InDels) specifically found in cell types from patient lines as indicated. Grouping of variants present in cell lines derived from 3 patients in Venn diagrams indicates absence of positive selection for any mutations. The majority of identified cell type specific variants is found in intergenic or intronic sequences (pie charts). No gene ontology (GO) term enrichment was found in identified variants. **(D)** Aligning all shared iPS cell/iSC-specific variants identified by k-means clustering (Fig. 4B, S7B) and AF-cut-off filtering (Fig. 4C) with the used seed sequence of sgRNA C4 within a 25bp search radius that must contain a NRG PAM-motif does not identify any significant homologies. The outlier (circles) with the most similarity to sgRNA C4 still exhibits 6 mismatches (see text for details). **(E)** TIDE analysis of sgRNA C4-mediated CAS9-cutting in CO2 patient fibroblasts at the *COL7A1* on- target (top) and the most likely *in silico* predicted off-target, i.e. *NOTCH1* (bottom). Measurements were taken 100bp-50bp upstream (i.e. internal Ctrl) and 10-60bp downstream of the predicted cut sites. **(F)** Plots of normalized WGS coverage 1000bp up-/downstream of the *COL7A1* on-target (top) and the *NOTCH1* off-target (bottom) from fibroblasts and thereof derived iPS cells as indicated. Note the sharp drop of coverage at the *COL7A1* locus in iPS cells due to a heterozygous deletion of 654bp (see Fig. 2D-G). **(G)** Summary of (Fig. 4E-F) for all *in silico* predicted exonic and intronic off-targets and the *COL7A1* on-target (left column). For TIDE analysis (middle column), noise-signals from internal controls were subtracted from values measured at cut sites (Fig. 4E). No InDels were found at any off-target in 4 clonal iPS cell lines from 3 patients (Fig. 4F). Heterozygous 1bp, 10bp (not shown) and 654bp big deletions were detected on *COL7A1* on-target alleles of patients with homozygous mutations (i.e. CO1 and CO2). **(H)** Variants identified via the STAMPv2 oncopanel from fibroblasts and thereof derived iPS cells and iSCs as indicated. Germline variants (green) are found in all 3 cell types. A heterozygous mutation of the androgen receptor (red) stems from clonal expansion of a fibroblast sub-population (3%) already carrying this lesion. Blue indicates variants identified by secondary methods (see text and Fig. S7F-H) for details.

Reassuringly, our second strategy based on AF/odds ratio cutoff-filtering identified >89% of the variants called by k-means clustering as shared iPS cell/iSC-specific (Fig. S7D). Unlike k-means clustering, AF/odds ratio cutoff-filtering identified variants unique to iPS cells or iSCs only, albeit in much smaller numbers than shared iPS cell/iSC-specific variants (Fig. 4C). Of note, almost all iPS cell-specific variants were present in other cell types with an AF>0, suggesting that these variants were not introduced *de novo* (Fig. S7E). In contrast, shared iPS cell/iSC- and iSC-specific variants were infrequent in other cell types suggesting some of them are *de-novo* or amplified from a rare (AF<5%) pre-existing variant (Fig. S7E). There was no overlap of these variants between patients (Fig. 4C). Most cell type-specific variants map to intronic or intergenic regions. Gene ontology (G.O.) term analysis of all variants located at loci with predicted functional properties (i.e. exons, splice sites, UTRs, and promotor/enhancer elements) did not yield any significant enrichment (Fig. 4C).

To investigate potential guide-dependent CAS9 off-target mutations, we searched a window of 25 bases around all shared iPS cell/iSC variants for PAM-like NRG-motifs that are combined with sequence similarities to the used sgRNA C4 ^44^. We did not observe any NRG motif adjacent to a 20-mer containing less than 6 mismatches compared to the sequence of sgRNA C4 (Fig. 4D). The cutting efficiency of CAS9 misguided by sgRNAs containing more than 3 mismatches is reported to be exceedingly low ^45, 46^. Accordingly, TIDE analysis of fibroblasts transfected with sgRNA C4/CAS9 RNPs did not detect any significant InDel formation at any *in silico* predicted exonic or intronic off target, whereas the *COL7A1* on target site was cut with over 87% efficiency (Fig. 4E and G). As TIDE can detect a maximum InDel size of 50bp, we also visualized InDels via plotting the normalized coverage of WGS reads obtained from unperturbed fibroblasts, thereof derived iPS cells and iSCs around all *in silico* predicted exonic, intronic, and intergenic off targets and the *COL7A1* on target. All previously identified CAS9- mediated deletions on the non-repaired *COL7A1* allele displayed the expected decreased read coverage (Fig. 4F, compare to Fig. 2G). No other InDels in proximity of potential off target cut sites were observed when searching a 1kb or 1Mb window (Fig. 4G and data not shown).

The functional consequences of detected variants are hard to predict. Their random nature and frequent pre-existence in the heterogeneous patient fibroblasts suggests that pathogenic effects caused by our manufacturing process are unlikely to materialize. To exclude, however, the *de novo* introduction or clonal expansion of variants in potentially cancer-promoting genes we performed the Clinical Laboratory Improvement Amendments (CLIA)-accredited, high-coverage sequencing Stanford Actionable Mutation Panel for Solid Tumors (STAMP) ^47^. iPS cell lines DEB125-1, DEB135-10(B), DEB135-24(B), CO1-131 and CO1-173 displayed no STAMP-hits other than germline variants already present in unperturbed parental fibroblasts (Figs. 4H, S5H). iPS cell lines CO2-65(B) and CO2-48(C) and their iSC progeny exhibited a heterozygous mutation in the androgen receptor (AR) that STAMP did not detect in fibroblasts (Fig. 4H).

Targeted ddPCR analysis of this mutation in parental fibroblasts revealed however an AF of 3% of this variant (which is below the 5% detection limit of STAMP), suggesting clonal expansion of a rare somatic mutation or even a pre-cancerous lesion of this patient. Accordingly, many other iPS cell lines derived from patient CO2 by the same production run did not harbor this AR mt (Fig. S7F-G). These results highlight the merit of the STAMP oncopanel to exclude rare cell products with potentially pathogenic mutations.

### *In vivo* efficacy and favorable safety profile of patient-derived *COL7A1*-corrected organotypic skin grafts

To test the functionality of the corrected organoid iSCs, we transplanted ITGA6-enriched iSC cultures on the back of immunocompromised nude mice. With 3-12 manufacturing runs per line, all five *COL7A1*-corrected patient iPS cell lines produced viable grafts and formed human skin *in vivo* (Fig. 5A,B). Graft success was determined by formation of stratified epidermis consisting of K14^+^ basal cells and K10/Involucrin^+^ upper stratified layers, and detection of human-specific collagen 7 in the basement membrane zone (Fig. 5B). Successful grafts were stable for at least 1 month after transplantation and comparable to primary human keratinocyte grafts ^8, 48, 49^, thereby providing a biologically and clinically relevant endpoint.

**Figure 5:**
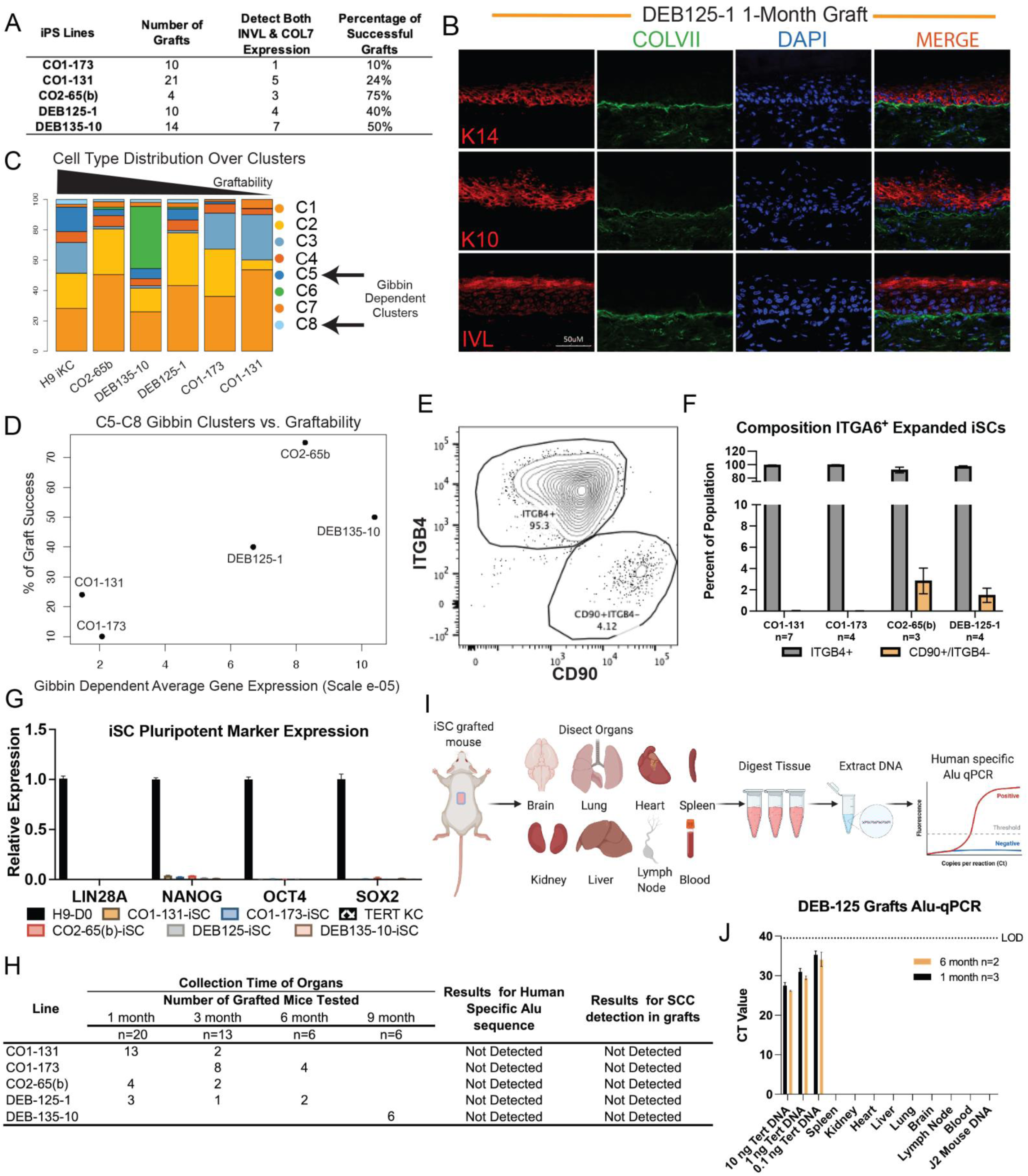
Patient iPS cell-derived orgonotypic skin grafts survive in mice with a favorable safety profile. **(A)** Table of mouse graft success determined by IF staining of basement marker Collagen 7 (COL7) and granular marker involucrin (IVL) at 1 month. Graft attempts include at least two distinct manufacturing runs **(B)** Representative IF image of 1 month DEB-125-1 derived iSC graft. Keratin (K)14, K10, involucrin (red), Collagen 7 (ColVII; green), DNA (blue). **(C)** Quantification (%) of 8 different cell type clusters (C1-8) comprising the DEBCT product and identified via individual scRNA sequencing in. **(D)** Strong positive correlation between % graft success rate in (Fig. 5A) and average Gibbin-dependent gene expression as quantified by scRNA sequencing. **(E)** Representative flow cytometry of cell composition of H9 ES cell derived iSCs labeled for ITGB4 and the Gibbin-dependent fibroblast marker CD90/Thy1 before grafting on to mice. **(F)** Bar graph of flow cytometry quantifying the average expanded iSC cell composition prior to grafting. **(G)** qRT-PCR detection of pluripotent marker expression (LIN28A, NANOG, OCT4 and SOX2) in the iSC product. H9 ES cells and TERT keratinocytes (KC) were used as controls. **(H)** Table summarizing Alu-qPCR-based biodistribution and histology-based tumor detection results from mice at 1-, 3- and 6-month post iSC grafting. **(I)** Diagram of method for detecting evading iSCs into mouse organs using human Alu-qPCR. **(J)** Representative Alu-qPCR from organs of DEB-125-1 derived iSC grafted mice at indicated time points. LOD is level of detectability in tissues spiked with human DNA from TERT keratinocytes.

While our manufacturing led to graftable iSCs from each line, we observed a line-dependent range of grafting efficacy (Fig. 5A). The different distribution of cell identity among the five lines characterized (Fig. 3K,L) enabled us to correlate the 4 epidermal, 3 dermal and 1 melanocyte cell groups of each line with its engraftment success rate. One prominent variable was the melanocyte cluster but that did not correlate with graftability (Fig. 5C, Fig. 3,KL). However, the lines with lowest grafting efficiency (CO1-173, CO1-131) distinctly lacked the two Gibbin- dependent fibroblast populations C5/C8, which were present in the other three lines (Fig. 3K,L, and 5C). Moreover, graftability correlated with the Gibbin-dependent dermal gene signature (Fig. 5D). To independently verify this finding, we measured the amount of Gibbin-dependent mesoderm in 4 iSC lines (n=3-7) by flow cytometry using the Gibbin-dependent marker CD90/Thy1 (Fig. S6I, Fig. 5E-F). Again, the two low efficiency lines exhibited few CD90+ cells whereas the better performing lines consistently produced 2-5% CD90+ dermal cells (Fig. 5E- F). These data suggest that optimal graftability requires the presence of Gibbin-dependent dermal cells, which have been shown to provide a critical maturation signal for epidermal stem cells ^31^.

The main safety concern of the iSC product is tumor formation. To assess potential teratoma formation from residual undifferentiated iPS cells, we first evaluated pluripotency marker expression in expanded ITGA6-enriched cells. *Lin28A*, *Nanog*, *Oct4*, and *Sox2* were below detectability in differentiation experiments of five different cell lines (Fig. 5G). Utilizing RNA expression of the pluripotency marker *Lin28A* we measured the limit of detection of contaminating iPS cells at 0.38% (i.e. 1 iPS cell in 260 immortalized keratinocytes ^50^; Fig S8A). In combination with absence of ITGB4 in iPS cells, which is necessary for adhesion to the BMZ ^34, 35^, we predicted the likelihood of teratoma formation to be low. Indeed, no teratomas were found in grafted animals. Another potential risk is the formation of squamous cell carcinoma (SCC), a common complication within the chronic wounds of adult EB patients. Careful investigation failed to show any histopathological signs of SCC formation in any of the human organoid skin grafts (Fig 5H). To assess the sensitivity to detect potential tumors in this assay, we performed spike-in positive control experiments using RDEB patient-derived SCC cells ^51^.

This defined the limit of detection (LOD) within the graft at 1 month at 0.025% SCC cells (Fig. S8B-C). To address the possibility of potentially metastasizing tumor cells from graft sites we designed a method to detect human DNA by Alu sequence qPCR from 7 organs and blood from all grafted mice at 1, 3 and 6 months (Fig. 5H-J). Spike-in experiments using 1-10,000 iSCs in 500K mouse cells from various organs demonstrated the LOD as 3,000 iSCs for lymph node, brain, and liver. Sensitivity for quantitation was determined to be 10,000 iSCs at 38 PCR cycles for all four organs tested (Fig. S8D). Example Alu-qPCR result for the DEB125-1 iSC grafted mice at 1 and 6 months showed no detection for human Alu sequences from the mouse organs or blood, with all iSC grafts tested. The 4 patient lines, grafted on mice, were all assayed for Alu-qPCR with no detection (Figs. 5H-J, S8E). To the level of detectability our data reinforces the safety and efficacy of our cell manufacturing method.

## DISCUSSION

With DEBCT manufacturing, we overcome existing technical hurdles to develop a scalable, GMP-compatible, and efficient platform for derivation of autologous and genetically corrected organotypic skin grafts for long-lasting treatment of RDEB patient wounds. This platform combines the generation and genetic correction of patient-derived iPS cells in a single manufacturing step and details a strategy for safe and reproducible manufacturing of graftable organotypic skin composites at clinical scale. The therapeutic product is composed of basal keratinocytes, dermal fibroblasts, and melanocytes, better resembling the composition of physiological tissues than previous approaches. Moreover, our use of generalizable manufacturing reagents and development of efficacy, toxicology and product characterization assays provide a reproducible regulatory and manufacturing path.

An important aspect of this study is the successful combination of CRISPR/CAS9-mediated gene editing and reprogramming into a single manufacturing step. Previous approaches generated iPS cells to be corrected in a subsequent clonal step, necessitating longer manufacturing time, higher cell expansion rates, and extensive additional quality control and release tests. Methodological simplification results in a series of advances: First, several new reagents enable clinical-scale corrected iPS cell manufacturing, including a novel transfection reagent that allows efficient delivery of three different nucleic acid/protein cargos with low toxicity, the high efficiency mRNA reprogramming kit, and the chemically defined iPS cell expansion media that ensures genomic/chromosomal stability. Second, the *COL7A1* correction and iPS cell reprogramming steps are seamless and lack genomic alterations to the DNA other than the corrected mutation and designed silent point mutations that facilitate genotyping.

Hence, our approach removes potential complications associated with most other gene and cell therapy approaches caused by random insertion mutagenesis, introduction of non-physiological gene regulatory elements, or alteration of endogenous loci. Third, genetically corrected autologous iPS cell banks can now rapidly be obtained and characterized in less than a month after dermal punch biopsy. This substantial acceleration not only increases the cells safety profile by lowering culture-induced mutation burden ^8^, but also reduces the time/cost of autologous iPS cell manufacturing, increasing the feasibility of this approach compared to off-the-shelf allogeneic iPS/ES cell-derived methods that suffer from host tissue tolerance issues ^52^.

We discovered one important caveat of CAS9-mediated editing. Whenever we observed integration of ssODN sequences on one allele, we almost always found InDel mutations on the other, which agrees with error-prone non homologous end-joining (NHEJ) being the predominant repair mechanism for DNA breaks in human cells ^53^. These InDels can be very large (Fig.2), resulting in standard PCR genotyping failing to amplify the affected allele, giving the false impression that genetic correction was bi-allelic. Our quantitative ddPCR analysis indicated that of 479 iPS cell lines derived from homozygous patients, only 3 exhibited potential bi-allelic integration of donor sequences (i.e. <1%; Fig. S4). This contrasts with reports of significantly higher bi-allelic *COL7A1* editing events in primary RDEB keratinocytes and iPS cells. Studies using non-quantitative PCR genotyping approaches may want to consider this potential problem. We suggest to carefully characterize the other allele and target the downstream of the two compound heterozygous mutations to minimize potential undesired consequences of InDel mutations ^54–60^.

Our cell manufacturing method for deriving engraftable organotypic skin equivalents is tailored for scalable GMP production in a 45-day process that is defined, xenofree, extensively validated ^61, 62^ and overcomes two key cell manufacturing hurdles. First, the induction of surface ectoderm, mesoderm and neuroectoderm mimics developmental signaling required for proper tissue maturation including development of epidermal, dermal, and melanocyte precursors. We find a significant proportion of the final iSC product to be the holoclone population previously identified as the long-term keratinocyte maintenance population ^41, 42^. In contrast to the previous culture intensive methods of subcloning and the use of a non-reproducible mouse fibroblast feeder line for maintenance, our inductive method produces all cell types necessary for an organotypic therapeutic product in scalable quantities via a defined and xenofree method. In addition, our studies support our previous results that underline the importance of epidermal-mesodermal signaling for subsequent induction of the region-dependent epidermal stratification program ^31^, as iPS cell clones deficient in differentiating into the Gibbin-dependent dermal population had a lower grafting efficacy. The reason for this line-to-line variability remains undetermined but the identification of the Gibbin-dependent CD90 surface marker, expressed on a small subpopulation of necessary mesodermal-like iSC cells, may provide a quantifiable biomarker predicting efficacy of graftability. Future advancements may include identification of distinct mesodermal signals that could give rise to distinct epidermal stratification programs, facilitating more subtle tissue morphologies with minimal morphogen reagent concentrations.

Second, incorporation of our iSC stem cell enrichment step overcomes maturation heterogeneity that reduces epidermal polarity and graft stratification. Despite a defined multi- lineage organoid culture aimed to regulate epidermal-dermal induction, immature cellular products appear to dominantly reduce the polarity and stratification abilities, requiring their removal for optimal grafting performance. While the differentiation efficiency varied between patient clones, our development of an ITGA6 enrichment step allowed derivation of organotypic skin grafts at clinical scale from all patients included in the study. Cell sorting technology for enriching antigen-specific cells is being used in various approved clinical products including CAR T cell and CD34+ cell enrichment ^63^. Because ITGA6 and its binding partner ITGB4 are present at elevated levels on most epithelial stem cells, our enrichment method will be applicable to other epithelial tissues.

While our abbreviated manufacturing process greatly reduced the number of cell doublings and consequently the chance of culture-induced mutations ^64^, our detailed characterization by whole and targeted genome sequencing paired with animal toxicology provides a rigorous assessment of manufacturing mutational and associated pathological risk. A key question was whether genetic variants in the therapeutic product are introduced during cell manufacturing or were pre-existing in patient skin cells. Sensitive ddPCR and WGS techniques confirmed that many variants stem from clonal amplification of pre-existing somatic mutations rather than *de novo* manufacturing-induced mutations. In agreement with high genomic stability within our manufacturing process, we found no variants unique to iPS cells only. We note that in line with naturally occurring proliferation-induced random mutagenesis ^64^, a small fraction of all detected variants does, however, arise *bona fide*, as indicated by rare iSC-specific variants that do not have any reads in parental fibroblasts and clonal parental iPS cell lines (Figs. 4C, S7). We did not observe any overlap of such iSC-specific variants between lines derived from different individuals or the same patient. This reflects the random nature of naturally occurring somatic mutations and legitimates our advancement that now allows DEBCT manufacturing in an unprecedented short time, thereby minimizing the risk of introducing a haphazard mutation with deleterious consequences. We did observe a three-fold line-to-line variability in the total number of SNVs arising in the different iPS cell lines (∼1,300 to ∼3,900). However, since many variants arise from clonal amplification of pre-existing somatic mutations, we conclude that the variation derives from the different mutational load of patient fibroblast.

Only three variants in only one detection method (k-means clustering) were found to be shared between patients and all of them were pre-existing. Similar results were found by the CLIA-certified STAMPv2 next generation sequencing assay to detect cancer-driving variants.

Corresponding to our sequencing analysis, sensitive toxicology assays for residual pluripotent or carcinogenic cellular components found no evidence for local or distant invasive properties of the product. This included grafts derived from patient cells harboring a mutation in the androgen receptor, which scored positive on the STAMPv2 screen. Given the greater ease and clinic-ready nature, we conclude that the STAMPv2 variant detection assay, along with our toxicology assays, will ensure a well-defined cellular product with low risk originating from our iPS cell-derived tissue manufacturing methods.

## ACKNOWLEDGEMENTS

We thank members of the Wernig and Oro labs for helpful discussions. Meredith Weglarz of the Stanford Shared FACS Facility for her expertise in cell purification and flow cytometry. This work was funded by generous grants from the EB Research Partnership (AEO, DR, and AC), the California Institute for Regenerative Medicine (TRAN1-10416, DISC2-12590), NIH (ARO73170), CIRM Scholars Program, Department of Defense (W81XWH-18-1-0706), and DEBRA Austria. K.R.R. and T.M.N were supported by R01GM121932. We are grateful to Dr. Dr. Andreas Reinisch (Medical University of Graz, Austria) for his help with identifying a suitable transfection reagent.

## MATERIALS AND METHODS

### Derivation and culture of primary patient fibroblasts

Fibroblasts from patients DEB125 and DEB135 were derived from a fresh dermal punch biopsy. Minced pieces (∼1mm^3^) of biopsies were cultured in DMEM (Gibco 12-430-062) with 10% fetal bovine serum (FBS; Hyclone SH30406.02 New Zealand sourced) and 1% Penicillin-Streptomycin (Fisher Scientific 15140163). Media was changed every 4 days and fibroblasts started to grow out from the tissue at day 4 and 8, respectively. Upon culture to confluency, fibroblasts were trypsinized (TrpLE; Gibco A1285901) and stored in Cryostore CS10 (Stem Cell Technologies 7930) in the vapor phase of liquid nitrogen. Fibroblasts from patients CO1 and CO2 were a generous gift from Dr. Dennis Roop (University of Colorado) and cultured in the above media.

### Single step editing/reprogramming

Immediately after transfection of patient fibroblasts with RNPs and ssONDs (see below), cultures were expanded to 225k cells in order to reprogram 3 wells of a 6 well dish. iPS cells were induced with a reprogramming kit a generous gift of iPEACE Inc (Los Altos, CA) according to the manufacturer’s recommendations. Briefly, 75k cells were seeded per well of a 6 well plate previously coated with iMatrix (Reprocell NP892-012) and transfected (VFS2205; see above) with 150ng of reprogramming mRNAs (iPEACE Inc) for 10 consecutive days. Cultures were subsequently maintained in StemFit media (Nacalai USA Basic03) according to the manufacturer’s recommendations (StemFit was supplemented with basic FGF from Preprotech AF-100-18B and Rock inhibitor from Axon 1683). After emergence of iPS cell colonies, non-reprogrammed cells encasing iPS cell colonies were removed as outlined in Fig. S3A-C and remaining iPS cells were incubated for an additional 24h before manual picking of colonies. Isolated iPS cell colonies were seeded in iMatrix-coated 48 well plates and maintained in StemFit media. After expansion of clonal lines, cultures were duplicated using the procedure recommended by Nacalai USA. gDNA was extracted from 1 duplicate for ddPCR analysis (see above) and sister cultures were maintained for further analysis.

### iPS and H9 ES cell differentiation

iPSC lines were maintained in StemFit^®^ Basic02 media (Anjinomoto), after expansion the cells were prepared for embryoid body formation using the AggreWell™ EB 400 microwell plate following the manufacturers recommendations (StemCell Technologies). Cells were dissociated in Accutase (Innovative Cell Technologies) counted and added at 1.2 million cells per microwell in AggreWell media containing 10 μM ROCK inhibitor (StemCell technologies 72302). Media was carefully changed the following day to remove ROCK inhibition. Embryoid bodies were collected after 48 hours and plated on 10 cm plates at roughly 250 Embryoid bodies per plate in StemFit^®^ Basic02 media until cell attachment on vitronectin (Gibco^TM^) pre coated plates following the manufactures recommendations. To initiate differentiation of PSCs to keratinocytes, the PSCs were induced with Essential 6 media (Gibco^TM^) containing 5 ng/mL BMP4 and 1 μM RA for 7 days; Defined Keratinocyte Serum Free Medium DKSFM (Gibco^TM^) keratinocyte selection media was continued through day 45 in differentiation and continued after MACS enrichment. The enriched keratinocytes were seeded onto Corning PureCoat™ ECM Mimetic 6-well Collagen I Peptide Plate (Corning). H9 ESC were maintained on matrigel hESC qualified matrix (Corning) and in Essential 8 media (Gibco^TM^) until differentiation began same as described above.

### ITGA6 enrichment by Miltenyi AutoMACS and CliniMACS separation

Differentiated day 45 cells were dissociated with Accutase (Innovative Cell Technologies) up to 30 minutes, washed, and counted in 10-20 mL of wash buffer containing PBS (Gibco), 1μM EDTA (Lonza), 2% BSA (Miltenyi) and 10 uM ROCK inhibitor (StemCell technologies 72302). For all wash steps, cells were pelleted in the wash buffer at 1000 rpm for 5 minutes. Cell pellets were resuspended in FcR Blocking reagent (Miltenyi), 20 μL FcR to 80μL of wash buffer up to 1×10^7^ cells for 5 minutes at room temperature. CD49F biotin antibody (REA518), was added to the blocked cells at 1:50 up for 1×10^7^ cells and incubated for 20 min at room temperature. After incubation, the wash buffer was adjusted to 10 mL and centrifuged at 1000 rpm for 5 minutes, then aspirated to completely leaving the cell pellet. Cells were resuspended in 80 μL of buffer and 20 μL of Anti-biotin IgG microbeads up to 1×10^7^ cells (Miltenyi Biotec) and incubated at room temperature for 15 minutes. Cells were then washed as previously described and resuspended in fresh wash buffer up to 4 mL. The labeled cell suspension was MACS separated using AutoMACS (Miltenyi Biotec) PosselD program setting. For CliniMACS cell enrichment the protocol was modified slightly, cells were labeled as described using the REA518 antibody, the Anti-biotin reagent (microbead) was determined by manufacturers recommendation at 1mL of microbead to 12.5 mL of CliniMACS buffer with 2% HSA, 1µM ROCK inhibitor. Program settings were optimized using; 1.1 (gentle), 5.1 (higher purity) and CD34.1 (lower purity, higher viability). Following separation, the cells are resuspended in Defined Keratinocyte Media (Gibco) and plated on ECM Collagen I coated peptide plates (Corning).

#### Production of RNPs

RNPs were generated by mixing sgRNAs with recombinant CAS9 at a molar ratio of 6:1, followed by incubation at room temperature for 30 minutes. Briefly, to generate 100ul with a concentration of 0.5 pmol RNP / ul we combined 81.13ul PBS with 18ul sgRNA (650ng/ul H2O) and 0.87ul CAS9. CAS9s were sourced from Integrated DNA Technologies (IDT; Alt-R® S.p. Cas9 Nuclease 1081058 or Alt-R® S.p. HiFi Cas9 Nuclease 1081060) and Aldevron (SpyFi Cas9 Nuclease 9214-0.25MG). sgRNAs were sourced from Synthego and had the following sequences fused to an 80-mer SpCas9 scaffold: C1 GGAUCCACCGUGAGUCCUCG, C2 GGGAUCCACCGUGAGUCCUC, C3 CGGGAUCCACCGUGAGUCCU, C4 ACUCACGGUGGAUCCCGCUG, C5 GACUCACGGUGGAUCCCGCU, C6 GGACUCACGGUGGAUCCCGC, and DEB135 ACUGGCACCAUCUCAACCUG.

#### ssONDs

ssONDs were sourced from IDT and stored as 10uM stock aliquots in H2O at -20C. The ssONDs used to edit the Colorado mt allele covered the genomic sequence of chromosome 3 between coordinates 48,570,186 – 48,569,987 (GRCh38.p13) and included the 4 silent mutations described in Fig1B. The ssONDs used to edit fibroblasts of patient DEB135 covered the genomic sequence of chromosome 3 (GRCh38.p13) between the following coordinates and included the 4 silent mutations described in Fig. S5A: ssOND 84bases - 48,572,943 – 48,572,860; ssOND 127bases - 48,573,004 – 48,572,878; ssOND 200bases - 48,572,943 – 48,572,744.

#### Transfection with RNPs and ssONDs

One day before transfection, 25k fibroblasts were seeded in penicillin-streptomycin-free media in a well of a 24-well plate. Transfection of 5pmol RNP (or of amounts indicated in Fig. S1D) and 10pmol ssOND was performed with VFS2205 transfection reagent (Vivofectamine™ Services from Thermo Fisher Scientific) according to the manufacturer’s recommendations.

#### Extraction of genomic DNA

Genomic (g)DNA was extracted using the Quick DNA miniprep kit (Zymo Research D3025) according to the manufacturer’s recommendations. Genomic DNA was eluted and stored in nuclease-free H2O (Ambion AM9937).

#### TIDE assay

TIDE assay ^65^ was performed as per the inventor’s instructions via the online tool at https://tide.nki.nl/. Edited loci were amplified from gDNA extracted 3 days (or 7 days for Fig. 4E and G) after transfection with RNPs and ssONDs, using the primers outlined in Table S1 and Q5 Hot-start high fidelity polymerase (New England Biolabs M0494S). PCR products were extracted from standard agarose gels using the QIAquick Gel Extraction Kit (Qiagen 28704) and sequenced by Elim Biopharm (www.elimbio.com).

#### ddPCR

gDNA was used as a template for droplet digital (dd) PCR according to manufacturer’s instructions for the BioRad QX200 system. ddPCR reactions were prepared using 2x ddPCR supermix (BioRad 1863025), bi-allelic reference-HEX primer/probe mix (primer 1: GGATGGGGAATGCAGCTCTT, primer 2: AGTGCGGCAGAATACAGCA, probe:5’HEX/TGATGGGTT/ZEN/GTGAAGGCAGCTGCACCT/3’IABkFQ), and one of the following FAM-conjugated primer/probe mixes: edited Colorado mt allele-FAM (primer 1: GAGTCAATGAACCTAATGTC, Primer 2: AGAGAGTCCTGGGGTA, probe: 5’6- FAM/AAGGGGGCT/ZEN/CACGGTG/3’IABkFQ), or edited DEB135 mt allele-FAM (primer 1: GACAGAGCTCTTCCCTCTCA, primer 2: CTGCCCCCAGAACACATAC, probe: 5’6- FAM/TGGCCGAGA/ZEN/CGGTGCC/3’IABkFQ). After generation of droplets in the BioRad QX200 generator, which employed DG8 cartridges (BioRad 1864008), gaskets (BioRad 1863009), and droplet generation oil for probes (BioRad 1863005), ddPCR mixes were loaded into 96 well plates (Fisher Scientific E951020346) and sealed with pierceable foil heat seal (BioRad 1814040). PCR reactions were run in a thermocycler using the following parameters: 1x 10 minutes at 95C, 47x 30 seconds at 94C followed by 1 minute 5 seconds at 57C, and 1x 10 minutes at 98C. Subsequently, ddPCR reactions were analyzed in a BioRad QX200 droplet reader and data analysis was performed with BioRad’s QuantaSoft Analysis Pro 1.0.596 software. The percentage of edited *COL7A1* alleles was calculated via the following formula: ((concentration in copies per ul of edited Colorado or DEB135 allele [FAM signal]*100) / concentration in copies per ul of bi-allelic reference [HEX signal])*2 = % heterozygously edited cells. For the competitive ddPCR assay detecting the wt and mt (c.1427G>A, p.G476E) allele of the androgen receptor (*AR*), following primer/probe mixes were employed: primer 1: GAAGGCCAGTTGTATGGAC, primer 2: CACATCAGGTGCGGTGAAG, AR-wt-FAM: 5’6- FAM/AGGCGGGAG/ZEN/CTGTAGCCC/3’IABkFQ, AR-mt-HEX: 5’HEX/AGGCGGAAG/ZEN/CTGTAGCCCC/3’IABkFQ. Competitive ddPCR assays were run as above with the following thermocycler parameters: 1x 10 minutes at 95C, 45x 30 seconds at 94C followed by 1 minute 5 seconds at 56C, and 1x 10 minutes at 98C. The percentage of *AR*mt alleles was calculated via the following formula: ((concentration in copies per ul of AR-mt [HEX signal]*100) / concentration in copies per ul of AR-mt [HEX] + AR-wt [FAM])*2 = % cells with *AR* mt. All ddPCR primer/probe mixes were sourced from IDT as PrimeTime Std qPCR Assays with a primer/probe ratio of 3.6.

#### Topo cloning

Topo cloning of PCR-amplified *COL7A1* alleles was achieved using the Zero Blunt TOPO PCR Cloning Kit, with One Shot TOP10 Chemically Competent E. coli cells (Thermo Fisher Scientific K2800-40) according to the manufacturer’s recommendations.

#### Sanger sequencing

Sanger sequencing was performed at Elim Biopharm according to the provider’s recommendations. Sequencing primers for Topo-cloned PCR products had following sequences: Fw 5’ GTAAAACGACGGCCAG ‘3, Rw 5’ CAGGAAACAGCTATGAC 3’. After purification from standard agarose gels (QIAquick Gel Extraction Kit, Qiagen 28704), PCR products were sequenced with either the Fw and/or Rw primer listed in Table S1.

#### *in silico* analysis of sgRNAs

sgRNA sequences were analyzed via the CRISPOR algorithm in order to identify activity and specificity scores and off targets ^66, 67^.

#### PCR

PCRs were performed with the Q5 Hot Start High-Fidelity 2X Master Mix (New England Biolabs M0494S) and primers outlined in Table S1. E. coli colony PCRs (Fig. 1F and Fig. S1F- H) were performed with CloneID 1X Colony PCR Mix (Lucigen 30059-2) and primers outlined in Table S1.

#### Immunofluorescence microscopy

Cells were fixed (4% paraformaldehyde), permeabilized (0.2% Triton X-100; except for Fig. 2H – TRA-1-81), blocked (BSA), and stained with antibodies against as per standard laboratory procedures. Nuclei were counterstained with DAPI. Images were acquired with a Leica DMi8 inverted microscope. Primary antibodies used: TRA-1-81 (Sigma Aldrich MAB4381), TRA-1-60 (Sigma Aldrich MAB4360), NANOG (Abcam ab21624), K18 (1:800, R&D AF7619), K14 (1:800, BioLegend SIG-3476-100), and p63 (1:100 Gene Tex GTX102425), ITGA6 (1:200, Millipore, MAB1378). Antibodies for mouse grafts comprised of rabbit anti-Involucrin (1:100, abcam), rabbit anti-keratin 14 (1:2000, Covance), mouse anti-keratin 18 (1:800 Abcam), rabbit anti-keratin 10 (1:500, Covance), human specific N terminal anti-collagen VII LH7.2 (1:250 Millipore), The fluorescence images were taken using the TCS SP5 confocal laser scanning microscope (Leica). Tissue sections were co-stained with 1:10000 Hoechst for 10 min. and slides were mounted with the Prolong Gold mounting medium (Life Technologies).

#### Karyotyping

Karyotyping was performed at WiCell or the Stanford University Medical Center Cytogenetics laboratory after expansion of iPS cells to at least 30 million cells.

#### Genomic DNA extraction and whole-genome sequencing

Genomic DNA (gDNA) was extracted from cell pellets of skin fibroblasts, iPS cells, and derived keratinocytes from three patients (11 total samples) with the MasterPure Complete DNA Purification Kit (Lucigen #MC85200). The gDNA yield was quantified by the Qubit dsDNA high-sensitivity (HS) fluorescence assay (Invitrogen # Q32851) and ranged from 39.6 to 106.0 ng/uL total in a final volume of 25 uL. The gDNA purity was verified by the NanoDrop 2000 spectrophotometer (Thermo Scientific #ND-2000). Tagmentation-based sequencing libraries were prepared from 500 ng gDNA in duplicate with the Illumina DNA prep (M) kit (Illumina #20018704) using IDT® for Illumina® DNA/RNA UD Indexes Set B, Tagmentation (Illumina #20027214). Libraries were sequenced in 150-bp paired end format on two sequential NovaSeq 6000 S4 lanes (Novogene Corporation Inc.). Each sample obtained at least 40X average coverage after combining the data from both library replicates.

#### Read pre-processing and variant calling

Raw reads were trimmed of adapter sequences using cutadapt^68^ in pair-end mode and subsequently mapped to the human hg38 reference genome (https://hgdownload.soe.ucsc.edu/goldenPath/hg38/bigZips/hg38.fa.gz) using BWA- MEM.^69^ The BAM files were processed by GATK4^70^ to sort mapped reads and mark PCR duplicates, followed by base quality score recalibration (BQSR) which utilized files of known human variant sites recommended in the best practice workflow of GATK4.

SNPs and/or Indels were called from processed BAM files using four variant-calling algorithms separately: HaplotypeCaller^70, 71^, Mutect2^72^, Lofreq2 ^73^ (SNP only) and Scalpel^74^ (Indel only). The overlapping hits of three SNP callers (HaplotypeCaller, Mutect2 and Lofreq2) and of three Indel callers (HaplotypeCaller, Mutect2 and Scalpel) were regarded as true variants. Functional annotations of the .vcf files were generated using ANNOVAR Wang, 2010, 20601685} with the embedded refGene protocol for hg38. Finally, only variants that do not match the dbSNP database^75^ (build 138) were considered as novel mutations and were included in subsequent analyses.

#### K-means classification and cell-type specific variant analysis

For each patient-derived lineage, novel mutations called in skin fibroblasts, iPS cells or derived keratinocytes were merged into one table with allele frequency (*AF*) information retained in all three cell types. Each variant *X* then has its *AF* information encoded as vector of three elements:

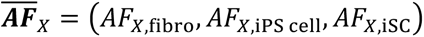

Unsupervised K-means clustering was conducted with all novel mutations using function “kmeans” in R package “stats”. Pseudo seeds were set for the random initiations of clustering centroids to ensure reproducible classification outcomes. Max iteration number was set as 1000 and “Lloyd” was selected as the clustering algorithm. The pre-defined cluster number K was tested from K=3 to K=9 in each individual task. The results of different patients consistently reveal a cluster exhibiting iPS cell/iSC-specific pattern at K=9 as highlighted in suppl Fig S7B. Therefore, we picked the smallest K numbers in each individual that defines such pattern (i.e. K=8 for CO1-131 and CO1-173; K=4 for CO2065(B); K=9 for DEB125-1), and generated variant list from the relevant clusters to be used in suppl Fig S7C.

To further identify variants that are specifically represented in iPS cell and/or iSC compared with their parental fibroblasts, we defined odds ratios *OR* to measure over-representation of a mutated allele in one cell type relative to another. For example, the odds ratio for variant *X* to be specifically represented in iPS cell but not the fibroblast is calculated as:

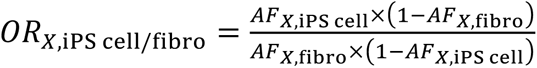

Any variants *X* passing the below filters were selected as iPS cell and/or iSC specific variants shown in Fig 4C:

(1) Shared iPS cell/iSC specific: (*AF_X,_*_fibro_ <0.25) *AND* (*AF_X,_*_iPS *cell*_ > 0.25) *AND* (*AF_X,_*_iSC_ > 0.25) *AND* (*OR_X,_*_iPS cell/fibro_ > 2.0) *AND* (*OR_X,_*_iSC/fibro_ > 2.0);
(2) iPS cell specific: (*AF_X,_*_fibro_ <0.25) *AND* (*AF_X,_*_iPS *cell*_ > 0.25) *AND* (*AF_X,_*_iSC_ > 0.25) *AND* (*OR_X,_*_iPS cell/fibro_ > 2.0) *AND* (*OR_X,_*_iPS cell/iSC_ > 2.0);
(3) iSC specific: (*AF_X,_*_fibro_ <0.25) *AND* (*AF_X,_*_iPS *cell*_ > 0.25) *AND* (*AF_X,_*_iSC_ > 0.25) *AND* (*OR_X,_*_iSc cell/fibro_ > 2.0) *AND* (*OR_X,_*_iSC/iPS cell_ > 2.0);

#### Coverage analysis of potential off-target sites

The 1KB up- and downstream regions of gRNA-targeted site (*COL7A1*) as well as 57 predicted off-target sites were investigated for potential alterations led by gRNA-dependent effects (Fig 4F). Normalized coverages were calculated as absolute read depth per site divided by the mean coverage of entire chromosome.

#### GO-enrichment analysis

GO enrichment analysis was performed via the online-tool at http://geneontology.org/ ^76, 77^.

#### STAMP analysis

The Stanford Actionable Mutation Panel of Solid Tumors (STAMP) version 2 sequencing was performed at Stanford University’s Anatomic Pathology and Clinical Laboratories.

### RNA-seq

Total RNA was isolated following Trizol reagent RNA extraction protocol for cultured cells. The RNA-seq libraries were constructed by TruSeq Stranded mRNA Library Prep kit (Illumina). All the libraries were sequenced to saturation on Illumina Hiseq2000 or NextSeq sequencers.

### RNA-seq Analysis

Fastq files were aligned to hg38 using TopHat 2.1.1 with parameters -p 10 --library-type fr-firststrand -r 100 --mate-std-dev 100. Aligned reads were processed to remove PCR duplicates using Samtools 1.8. Raw counts and RPKM values were calculated using HOMER analyzeRepeats.pl. To test for differential expression, raw reads were compared using DESEQ2, and filtered based on an adjusted *p-value* of < 0.05 and 2-fold change.

### qRT-PCR

Total RNA was isolated following Trizol reagent RNA extraction protocol for cultured cells. qRT-PCR was performed as described by TaqMan^TM^ RNA-to-CT^TM^ 1-step kit (Applied Biosystems). TaqMan probes available in reagent list.

### scRNA sequencing

Differentiated day 7 or enriched and expanded D50 iSCs were dissociated with Accutase (Innovative Cell Technologies) up to 30 minutes, filtered with a 40μm mesh, washed in Wash buffer previously described. Dissociated cells were counted and assayed with trypan blue for live cell counts. Collections with greater than 10% dead cells were processed for dead cell removal using a Dead Cell Removal kit (Miltenyi) following the manufacturers protocol. A total of 10,000 cells were resuspended in Wash buffer at 1,000 cells per μL. Library preparation was carried out following the Chromium Single Cell Chromium Next GEM Single Cell 3ʹ Reagent Kits v3.1 protocol.

### Single-cell RNA-seq analysis

FASTQ files were processed using 10x Genomics Cell Ranger v.3.1.0 and the human genome GRCh38. Cells with UMI counts greater than 500 and with mitochondrial percentage below 20% were included for further analysis. Downstream analyses were performed using Seurat v.4.0.0. 5 iKC samples and a H9KC sample were merged into one object and normalized using the default parameters. To compare between each clone, the Seurat object was split by samples and anchors were identified between samples using FindIntegrationAnchors, followed by integration. A total of 2,000 highly variable features were identified, objects were scaled to regress out cell cycle stages. Cells were clustered using 20 dimensions and a resolution of 0.2 to obtain 8 clusters. FindAllMarkers using log(fold-change) > 0.2 was used for differential expression analysis within individual clusters. For gene scoring, we used Gibbin dependent gene list from Collier A et al and Holoclone geneset from Enzo E et al to run the AddModuleScore function.

### Fluorescence activated cell sorting and Flow cytometry

Dissociated cells were washed with FACS buffer (2% BSA Cat# 130-091-376/1µM ROCK inhibitor/ AutoMACS rinsing solution Cat# 130-091-222). After wash steps, cells were fixed and permeabilized with (eBioscience™ Intracellular Fixation & Permeabilization Buffer Set, Cat # 88- 8824-00) then stained for antibodies of interest for 30 min at 4°C (FITC anti-K14, CBL197F from Millipore; PerCP anti-K18 (NB120- 7797, Novus), ITGA6 (PE anti-CD49F BD Cat #555736), all at 1:100 dilution. Enriched iSCs were analyzed using Streptavidin Alexa Fluor™ 647 conjugate (Life Technologies Cat # S32357, 1:500 dilution) and FITC-Labeling Reagent (Miltenyi Biotec, Cat # 327806, 1:50 dilution). Composition analysis of enriched iSCs was performed using anti- ITGB4 (BD Cat # 744150, 1:100 dilution) and anti-CD90 (BD Cat # 555595, 1:100 dilution).

Cells were strained through 35μm mesh. Flow cytometry data was acquired on a BD LSRII in the Stanford Shared FACS Facility with BD FACSDiva Software. Ten thousand events were collected. Analysis was performed with FlowJo software.

### *In vitro* skin reconstitution assay

Generation of organotypic epidermis was performed by following the protocol described previously ^8^ with minor modifications. Derived induced keratinocytes and unsorted passaged differentiated cells were expanded in Defined Keratinocyte SFM (Gibco^TM^) until confluent then passaged onto devitalized human dermis seeded at a density of 1×10^6^ cells. The medium was then gradually changed to 7F stratification media for 7 days, after which the dermal sheet was raised to the air-liquid interface. After 2 weeks, the reconstituted epidermis was collected for IF staining.

### Mouse skin engraftment with iPS cell derived iSC and Luciferase-RDEB-SCC lines

All animal experiments followed the NIH (National Institutes of Health) *Guide for the Care and Use of Laboratory Animals* under Stanford APLAC (Administrative Panel on Laboratory Animal Care). Xenograft protocol was performed as described previously ^8, 48^. ITGA6-enriched cells (1×10^6^) were seeded onto a 1.5-cm^2^ piece of devitalized human dermis (New York Firefighter Skin Bank) and grown in DKSFM (Gibco^TM^) for 10 days, followed by Keratinocyte Growth Medium (Gibco^TM^) for 5 days. Next, the pieces were grafted onto the backs of NSG mice for 1-6 months. Upon collection, the pieces were embedded in OCT and paraffin for immunofluorescence analysis. In vivo tumorigenicity of the reprogrammed and re-differentiated EB-SCC-iSCs expressing the luciferase gene, were created as previously described ^51^ were spiked with normal human keratinocytes NHKs to generate the 3D skin constructs and then grafted onto immunodeficient mice. Tumors were formed in mice grafted with RDEB-SCC-iSCs within 4 weeks and detected with the bioluminescence signal intensities.

### Alu-qPCR

In grafted mice, detection of emanating cells using Alu-sequence was performed on mouse organs including lymph nodes, liver, lungs, spleen, kidneys, heart, brain, and blood. The organs were resected and transferred to a sterile petri dish containing cold PBS, washed, finely minced, and weighed. DNA was extracted from the minced tissue according to the QIAamp DNA mini kit protocol (Qiagen Cat# 51304) with an overnight cell lysis at 56°C and 10-minute lysis for 100 uL blood. Alu-qPCR was performed using 10ng of extracted DNA in triplicate using amplification protocol by Applied biosystems (Universal Master Mix II, no UNG: Applied Biosystems Cat# 4440040 and Alu Probe: ThermoFisher Scientific Cat# 4351372).

## SUPPLEMENTARY FIGURES AND LEGENDS

**Figure S1:**
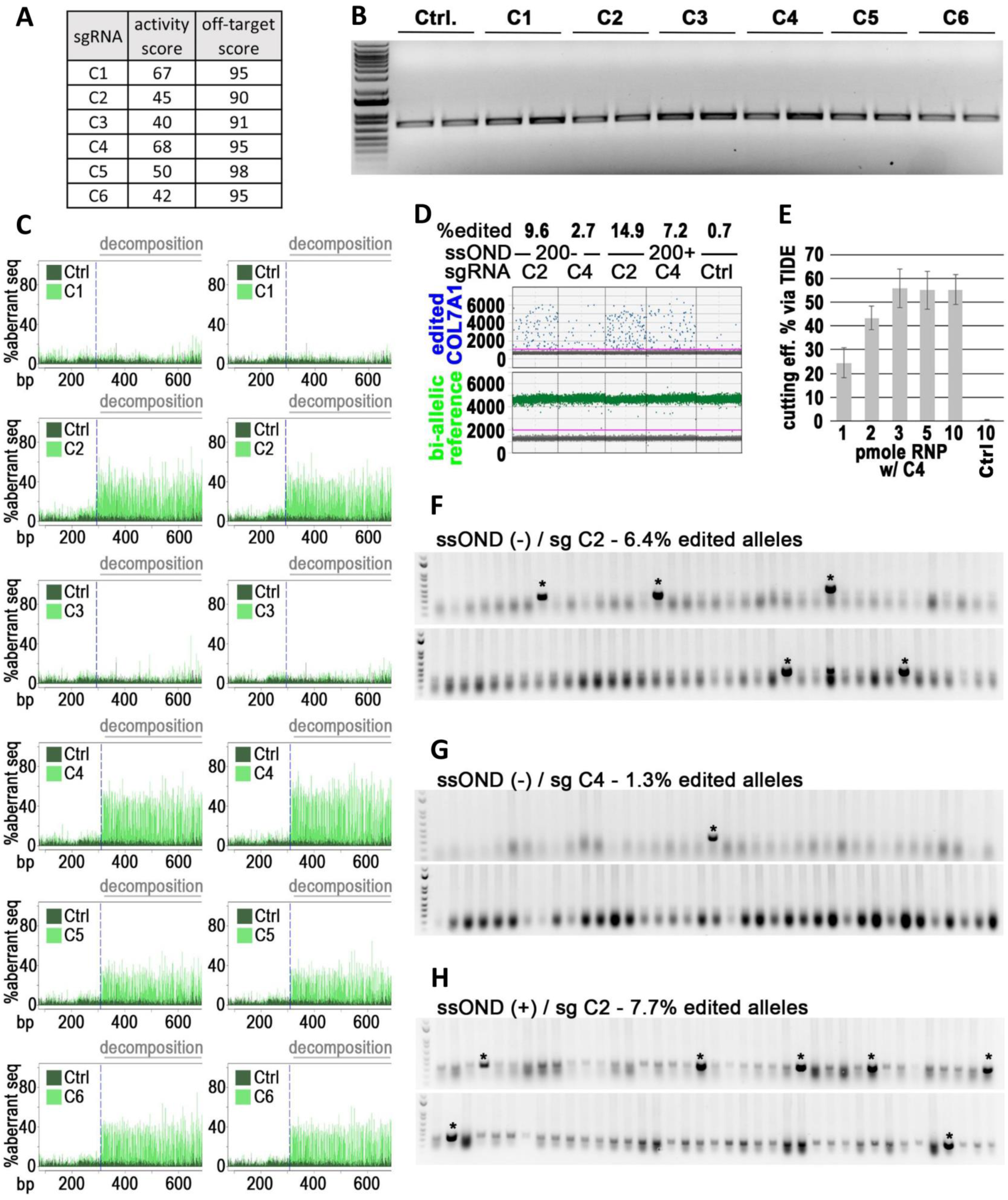
Optimized editing of the Colorado mt (7485+5 G>A). **(A)** *In silico* predicted efficiency (middle column) and specificity (right column) scores ^66, 67^ of the 6 tested sgRNAs (left column) used to mediate CAS9-cutting of the pathogenic Colorado allele. **(B)** Agarose gel visualizing PCR amplicons of a 731bp sequence surrounding the *COL7A1* target locus from homozygous CO2 patient fibroblasts transfected with RNPs containing CAS9 and indicated sgRNAs (Ctrl. omitted sgRNA). Replicate experiments are shown. **(C)** TIDE traces of PCR amplicons shown in (Fig. S1B). **(D)** *COL7A1* editing efficiencies as measured by ddPCR in DEB125 primary patient fibroblasts (het. Colorado mt) after transfection with ssODNs and sgRNA/CAS9-containing RNPs as indicated. A bi-allelic locus (green) is used as a reference for calculating *COL7A1* editing (blue) efficiencies. Ctrl omitted sgRNAs. **(E)** InDel formation in CO2 fibroblasts after transfection with indicated amounts of sgRNA C4-containing RNPs as measured by TIDE (n=2; stdev). Ctrl. omitted sgRNA. **(F-H)** Agarose gels visualizing 78 E. coli colony-PCRs to detect edited TOPO-cloned *COL7A1* alleles from CO2 fibroblasts transfected with ssODNs and RNPs containing sgRNAs as indicated. A primer specific for silent mutations (Fig. 1B) only amplifies PCR products of alleles with integration of donor sequences (asterisks).

**Figure S2:**
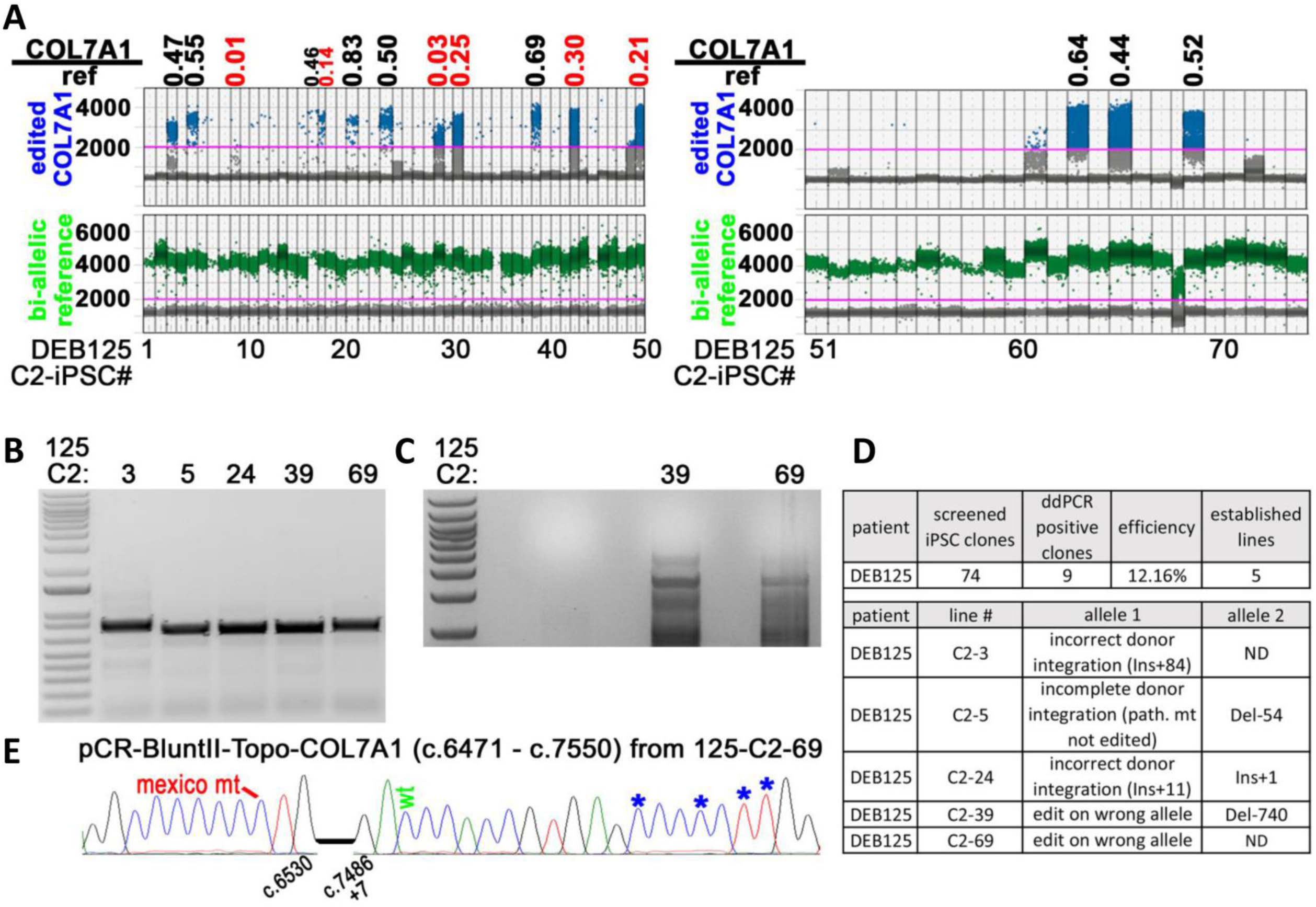
Single manufacturing step/editing reprogramming of patient DEB125 with the less specific sgRNA C2. **(A)** We assessed combined repair and reprogramming using sgRNA C2 and (+)ssODN in fibroblasts from patient DEB125. RNPs and the ssODN were transfected, followed by 10 consecutive daily doses of reprogramming mRNAs. This treatment yielded 74 iPS cell colonies. Screening via ddPCR, identified 9 candidate lines (∼12%) with potential mono- or biallelic integration of ssODN sequences at the *COL7A1* locus. This is consistent with the expected ∼15% *COL7A1*-editing frequency (Fig. S1D). Ratios of signals detecting edited *COL7A1* alleles (blue) and a bi-allelic reference locus (green) are used to identify mono-allelic (0.5 +/-0.19) or bi-allelic (1.0 +/-0.19) editing event (black values; red values below/above cut off indicate potentially mixed or incorrectly edited clones). **(B-C)** We established 5 of these lines and used their genomic DNA to further characterize the edited *COL7A1* locus. Agarose gel visualizing PCR amplicons of a 731bp (Fig. S2B) and 4560bp (Fig. S2C) sequence surrounding the edited *COL7A1* locus from established single-step edited/reprogrammed iPS cell lines from (Fig. S2A). **(D-E)** Cloning of the PCR amplified region followed by Sanger sequencing revealed that 2 of these 5 lines had correctly edited *COL7A1* alleles, displaying all intended silent mutations and a wt sequence at the Colorado mutation site. Shown is the summary of single-step editing/reprogramming screen conducted with sgRNA C2 and (+)ssODN from patient DEB125 (top). Topo-cloning and sanger sequencing of PCR products from (Fig. S2B-C) identified 2 iPS cell lines with correct donor integration (i.e. 125-C2-39 and 125-C2-69) that however, occurred on the wrong allele carrying the second heterozygous compound mutation of this patient (i.e. Mexico mt; 6527dupC). This is in agreement with the low specificity of sgRNA C2 for the Colorado-allele (Fig. 1D). We note that sgRNA C2 is capable of targeting the disease allele as we have successfully used it to derive such cell lines from fibroblasts of patient CO2 (homozygous for the Colorado mt; Fig. 4A).

**Figure S3:**
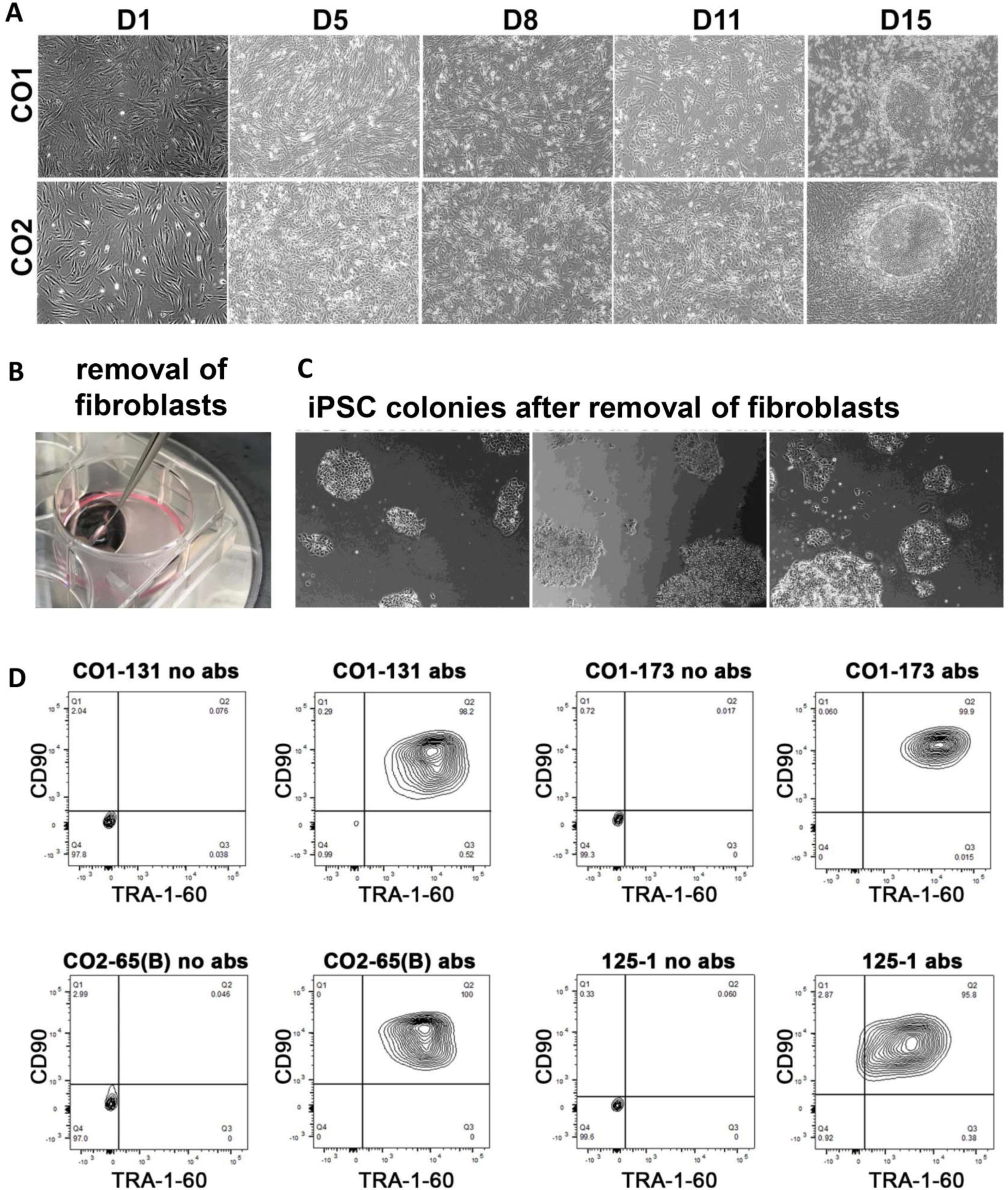
Efficient derivation of high-quality iPS cells from primary RDEB patient fibroblasts. **(A)** Phase contrast microscopy pictures of cell cultures during single-step editing/reprogramming from time points (D: day) and patients as indicated. iPS cell colonies emerge around D11-14. **(B)** Non-reprogrammed cells grow to confluency, adhere together, overlay iPS cell colonies and can be mechanically removed. **(C)** iPS cell colonies remain after removal of non-reprogrammed cells. **(D)** Flow cytometry analysis of single-step edited/reprogrammed iPS cell lines with and without labeling (antibodies; abs) of the pluripotency surface marker TRA-1-60 as indicated.

**Figure S4:**
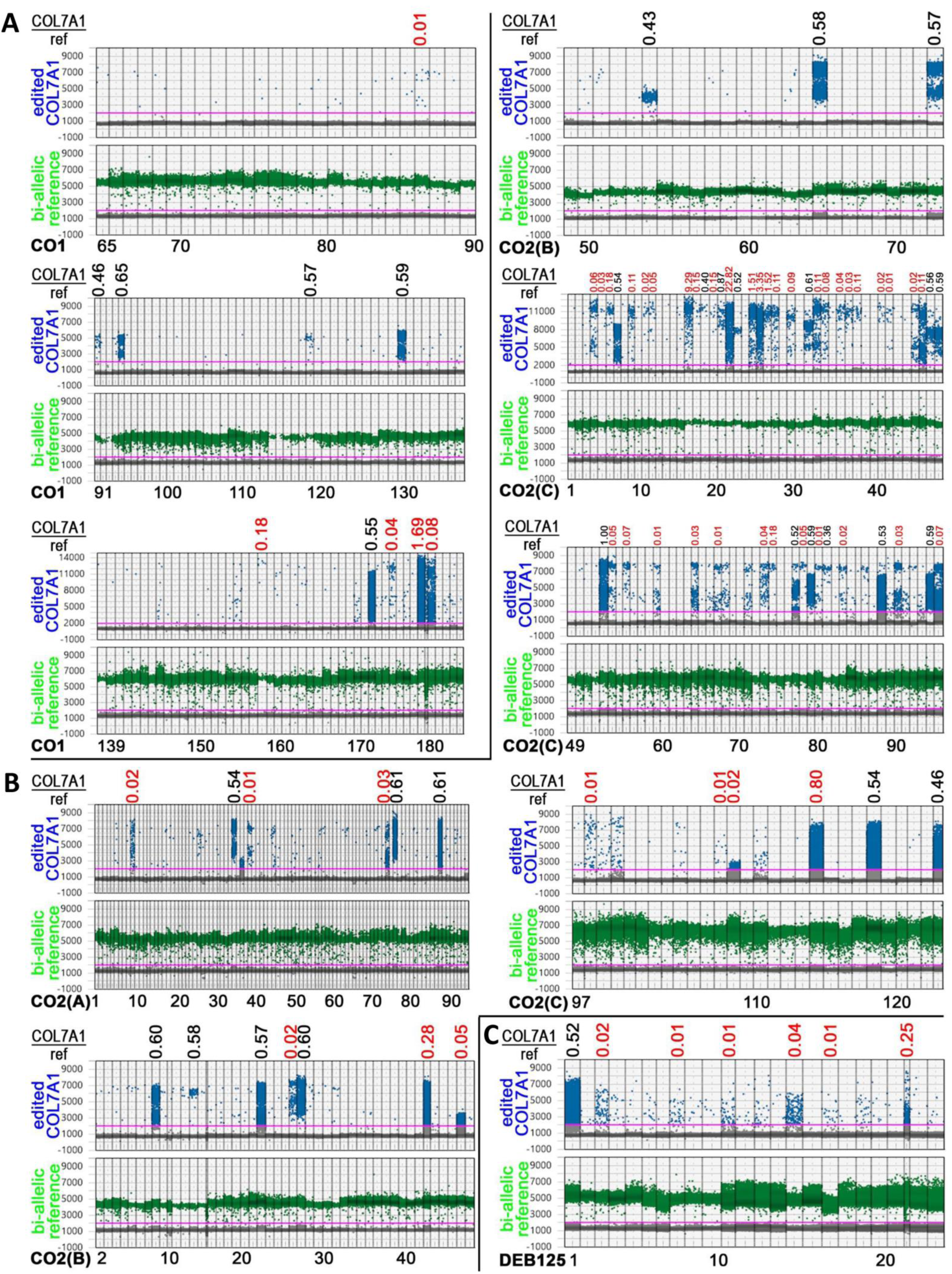
ddPCR screening of single-step edited/reprogrammed iPS cells. **(A-C)** Screen of 122 (A), 293 (B), and 24 (C) single-step edited/reprogrammed iPS cell lines via ddPCR from patients as indicated. Ratios of signals detecting edited *COL7A1* alleles (blue) and a bi-allelic reference locus (green) are used to identify mono-allelic (0.5 +/-0.19) or bi-allelic (1.0 +/-0.19) editing events (black values; red values below/above cut off indicate potentially mixed or incorrectly edited clones).

**Figure S5:**
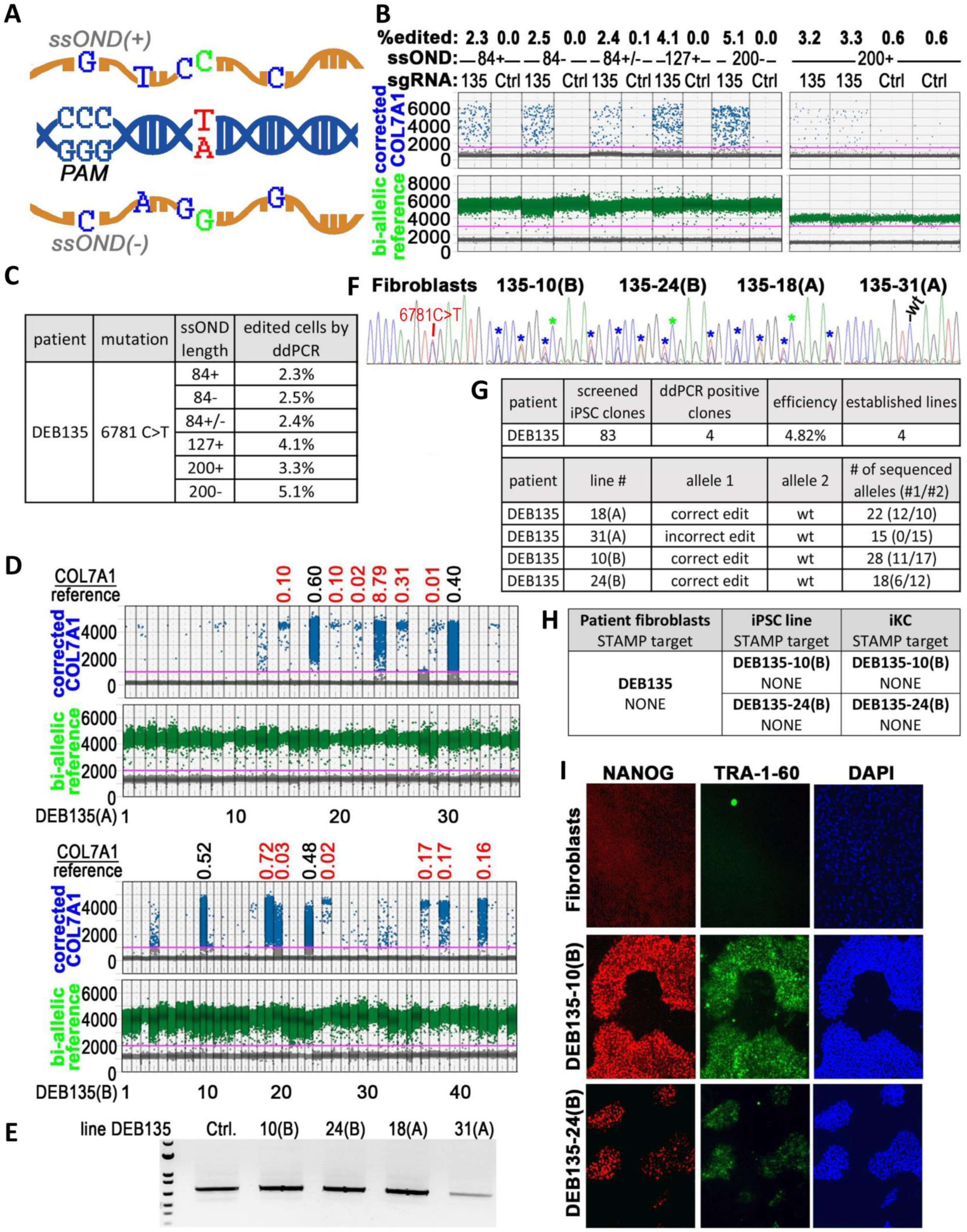
Optimized single manufacturing step editing/reprogramming of patient DEB135 (6781C>T). **(A)** Overview of the *COL7A1* target mutation 6781 C>T (red) and (+)/(-) ssODNs used for editing. PAM site to mediate cutting of pathogenic allele via CRISPR/CAS9 is shown. ssODNs encode for the wild type sequence (green) and 4 silent mutations (blue) used for detection of editing events. **(B-C)** *COL7A1* 6781C>T editing efficiencies as measured by ddPCR in DEB135 primary patient fibroblasts after transfection with ssODNs of various lengths, strandedness, and sgRNA 135/CAS9-containing RNPs as indicated. A bi-allelic locus (green) is used as a reference for calculating *COL7A1* editing (blue) efficiencies, assuming mono-allelic integration of donor sequence. The length of ssODNs correlates with editing efficiencies. Ctrls omitted sgRNAs. **(D)** Screen of 83 iPS cell lines via ddPCR after single-step editing/reprogramming of fibroblasts from patient DEB135 with (-)ssODN and sgRNA 135. Ratios of signals detecting edited *COL7A1* alleles (blue) and a bi-allelic reference locus (green) are used to identify mono-allelic (0.5 +/-0.19) or bi-allelic (1.0 +/-0.19) edited iPS cell lines (black values; red values below/above cut off indicate potentially mixed or incorrectly edited clones). **(E)** Agarose gel visualizing PCR amplicons of a 696bp sequence surrounding the edited *COL7A1* locus from established iPS cell lines from (Fig. S5D). Unedited fibroblasts were used as Ctrl. **(F)** Sanger sequencing traces of PCR amplicons shown in (Fig. S5E) containing both *COL7A1* alleles. Unedited fibroblasts show the heterozygous 6781C>T mutation as a double peak. Correctly edited iPS cell lines show heterozygous integration of intended silent mutations (blue asterisks) and repair of the pathogenic mutation (green asterisks). iPS cell line 135-31(A) only displayed the wild type sequence, indicative of a large InDel prohibiting amplification of the target allele. **(G)** Summary of single-step editing/reprogramming screen of patient DEB135 fibroblasts (top). Sanger sequencing of individual Topo-cloned alleles (Fig. S5E) confirmed correct editing in 3 iPS cell lines. See text for details. **(H)** STAMPv2 oncopanel sequencing of indicated cell lines did not identify any variants. **(I)** Immuno-fluorescence microscopy images of DEB135 fibroblasts and thereof derived iPS cells. NANOG (red), TRA-1-60 (green), and DNA (blue).

**Figure S6:**
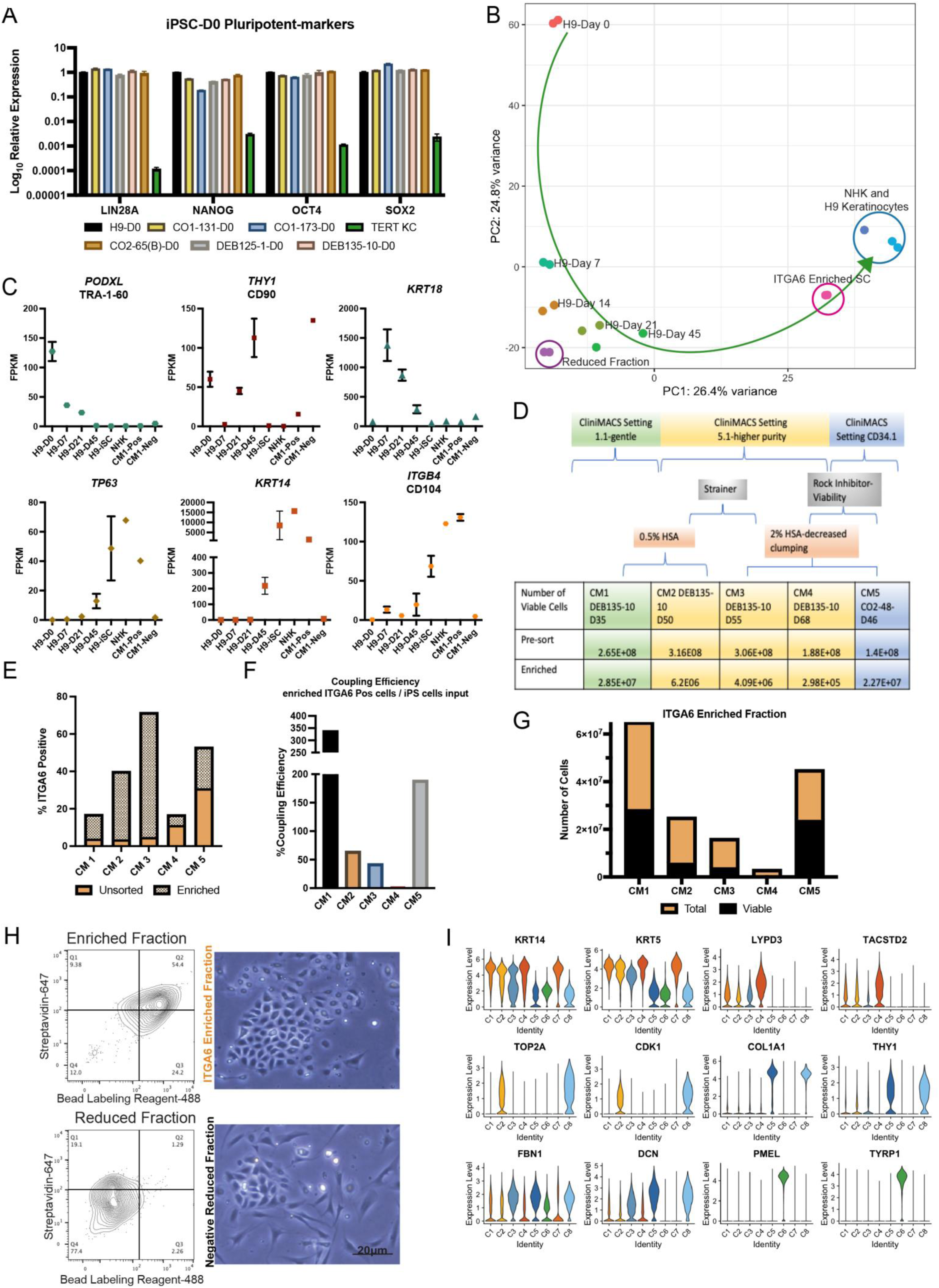
Efficient ITGA6 enrichment using CliniMACS purification for clinical scale manufacturing. **(A)** qRT-PCR of pluripotency marker expression of iPS cell lines at Day 0. TERT keratinocytes (KC) were used as the negative control. **(B)** Principal component analysis of RNA-seq from the CliniMACS ITGA6-enriched and reduced iSC cells compared to a H9-ES cells differentiation time course (D0-D45); H9 keratinocytes (KC) and NHK (positive control). **(C)** RNA-seq expression of CliniMACs (CM) enriched and reduced populations compared to H9 Day 0-45 differentiations, H9 iSC cells, and NHKs, illustrating the non-target population markers (TRA-1-60, Thy1/CD90, KRT18) and the target population markers (P63, KRT14 and ITGB4). **(D)** Table of optimizations for CliniMACs including program and reagents changes for each CM run with pre and post enrichment cell counts. **(E)** % ITGA6 positive cells of CM-sorted populations compared to the unsorted population by flow cytometry. **(F)** %Coupling efficiency (CE) determined by the equation %CE= live sorted iSCs/iPS cell input x 100 for CliniMACS runs CM 1-5. Note efficiency of CD34.1 automation program (i.e. CM 5). **(G)** Number of both viable and non-viable CliniMACS enriched iSCs. **(H)** Flow cytometry plots and corresponding bright field images of iSC Enriched and Reduced Fractions from CM3 CliniMACS separation (10x Magnification). **(I)** Violin plots of relative expression levels of key marker genes representing each cell cluster contained in the iSC product (Fig. 3), including basal keratinocytes (C1), cycling holoclones (C2), early differentiating keratinocytes (C4), Gibbin-dependent mesoderm (C5, C8), pre-vascular mesoderm (C3), and melanocytes (C6).

**Figure S7:**
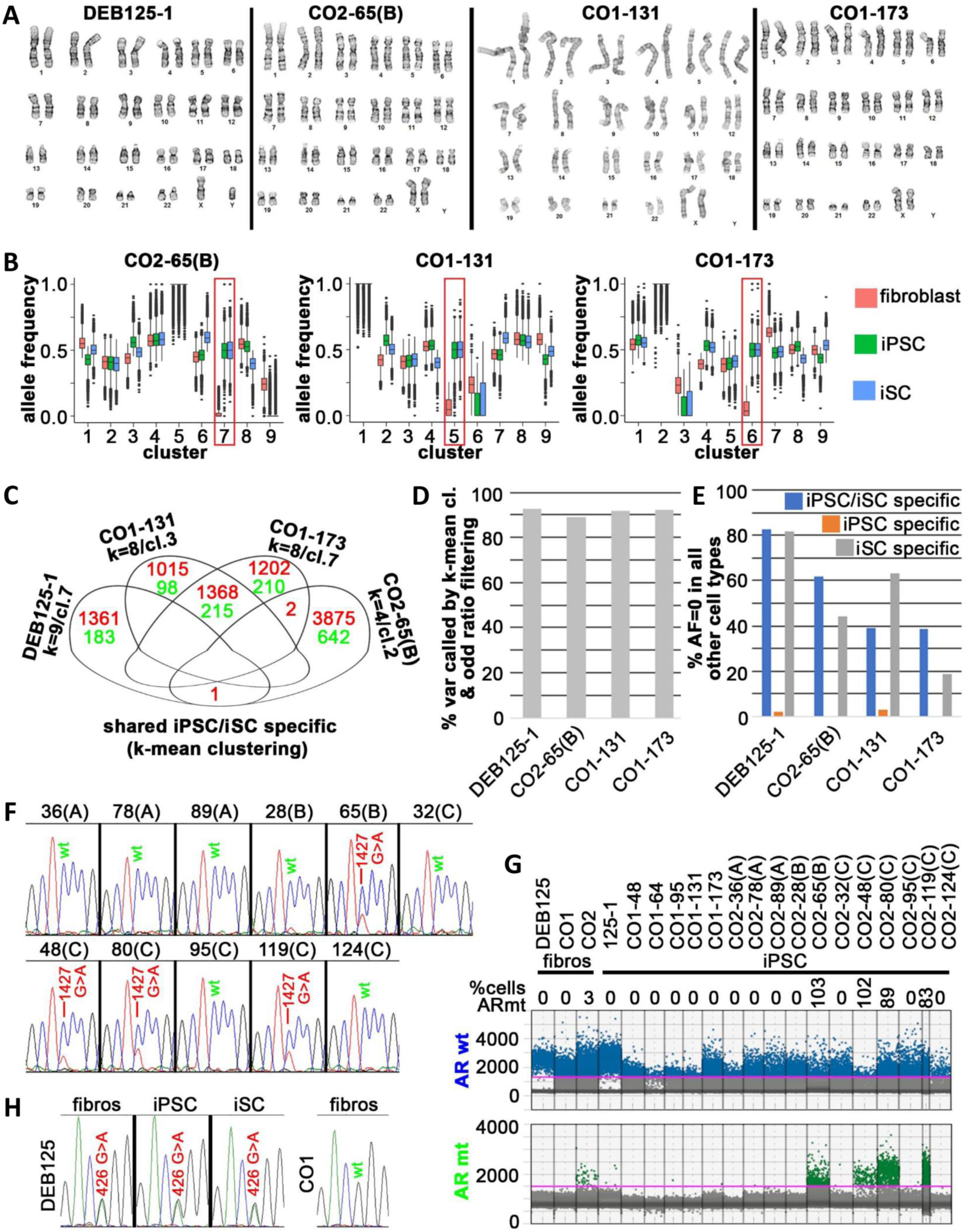
Genomic and chromosomal stability of single step edited/reprogrammed iPS cells and iSCs. **(A)** Representative normal karyotypes of four single-step edited/reprogrammed iPS cell lines from 3 individuals. **(B)** K-means clustering of all novel variants found by 40x whole genome sequencing (WGS) in fibroblasts and thereof derived iPS and iSC cells from patients CO1 and CO2 (CO2-65(B) n=114594; CO1-131 n=102278; CO1-173 n=101915). The red frame highlights differentially expressed allele frequencies (AFs) of variants in iPS/iSC cells compared to parental fibroblasts. **(C)** Variants (red: SNPs; green: InDels) that are specifically found in iPS and iSC cells of indicated cell lines as identified by k-means clustering. Grouping of variants in Venn diagrams indicates virtually no selection for mutations. Variants of clusters (cl.) showing clear separation between AFs found in fibroblasts and AFs found in iPS/iSC cells from feature spaces with the lowest k (indicated) were selected for this analysis. **(D)** AF cut-off filtering and k-means clustering identify virtually the same (>89%) variants specific to iPS/iSC cells. **(E)** Percentage of identified cell type-specific variants with no reads in other cell types as indicated. Note that virtually all iPS cell-specific variants identified by AF cut-off filtering (Fig.4C) were also found at lower AFs in other cell types. **(F)** Sanger sequencing traces of a PCR amplicon containing the androgen receptor (AR) locus c.1427 from indicated iPS cell lines of patient CO2. A heterozygous mutation was found in 4 of 11 iPS cell lines (double peaks). **(G)** A competitive ddPCR assay with probes specific for the wt and mt AR sequence from indicated cell lines confirms that 4 lines derived from patient CO2 harbor the mutation. Note that ddPCR detects this mtAR also in CO2 fibroblasts at low frequencies. **(H)** Sanger sequencing traces of a PCR amplicon containing the CDKN1B locus c.426 confirm a heterozygous germline mt (double peak) present in fibroblasts and thereof derived iPS/iSC cells of patient DEB125. Fibroblasts of patient CO1 were used as a negative control.

**Figure S8:**
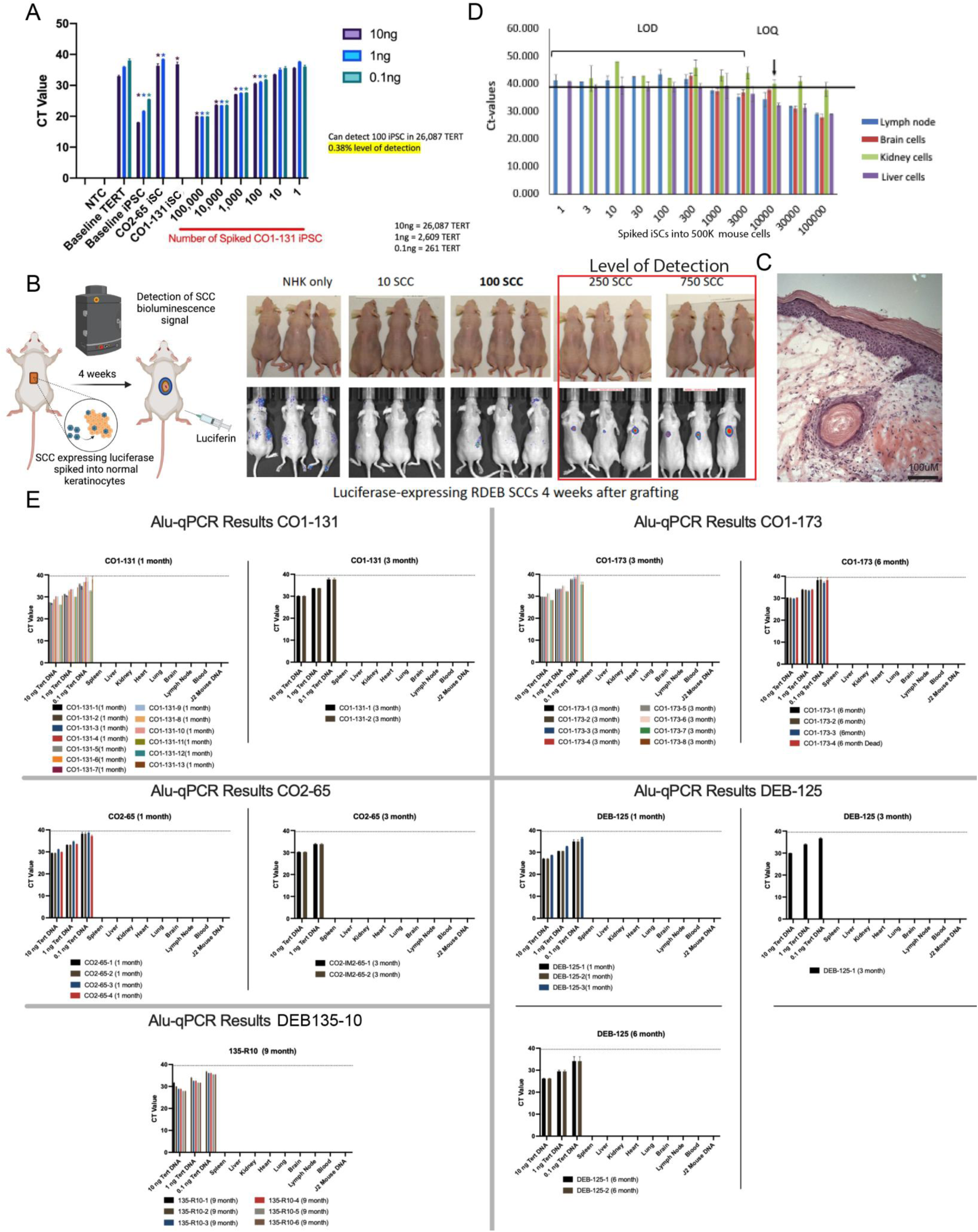
iPS cell detection assay, and qualification of human-specific Alu-PCR and tumor detection assays. **(A)** qRT-PCR of LIN28A to detect remnant iPS cells in expanded ITGA6-enriched cultures. **(B)** Schematic of assay to detect luciferase expressing-RDEB- squamous cell carcinoma (SCC)-cells in iSC cell grafts from a 4-week mouse tumor model. Bioluminescent detection of NHKs alone, 10, 100, 250 and 750 RDEB patient-derived SCC cells in 4-week-old grafts on mice to determine a level of detection (LOD) and **(C)** corresponding H&E histology of graft site. **(D)** Alu-qPCR detecting TERT-human keratinocytes in mouse cells from indicated organs to determine the level of detection (LOD) and quantification (LOQ) of human DNA for the biodistribution assay (SEM). **(E)** Alu-qPCR experiments performed on organs and blood of iSC grafted mice. DEBCT product from iPS cell clones as indicated and the time on the mouse (1-9 months). At left of each graph are the positive controls, with the tissue sampled on the X axis. Error bars are +/- SEM.

**Table S1:**
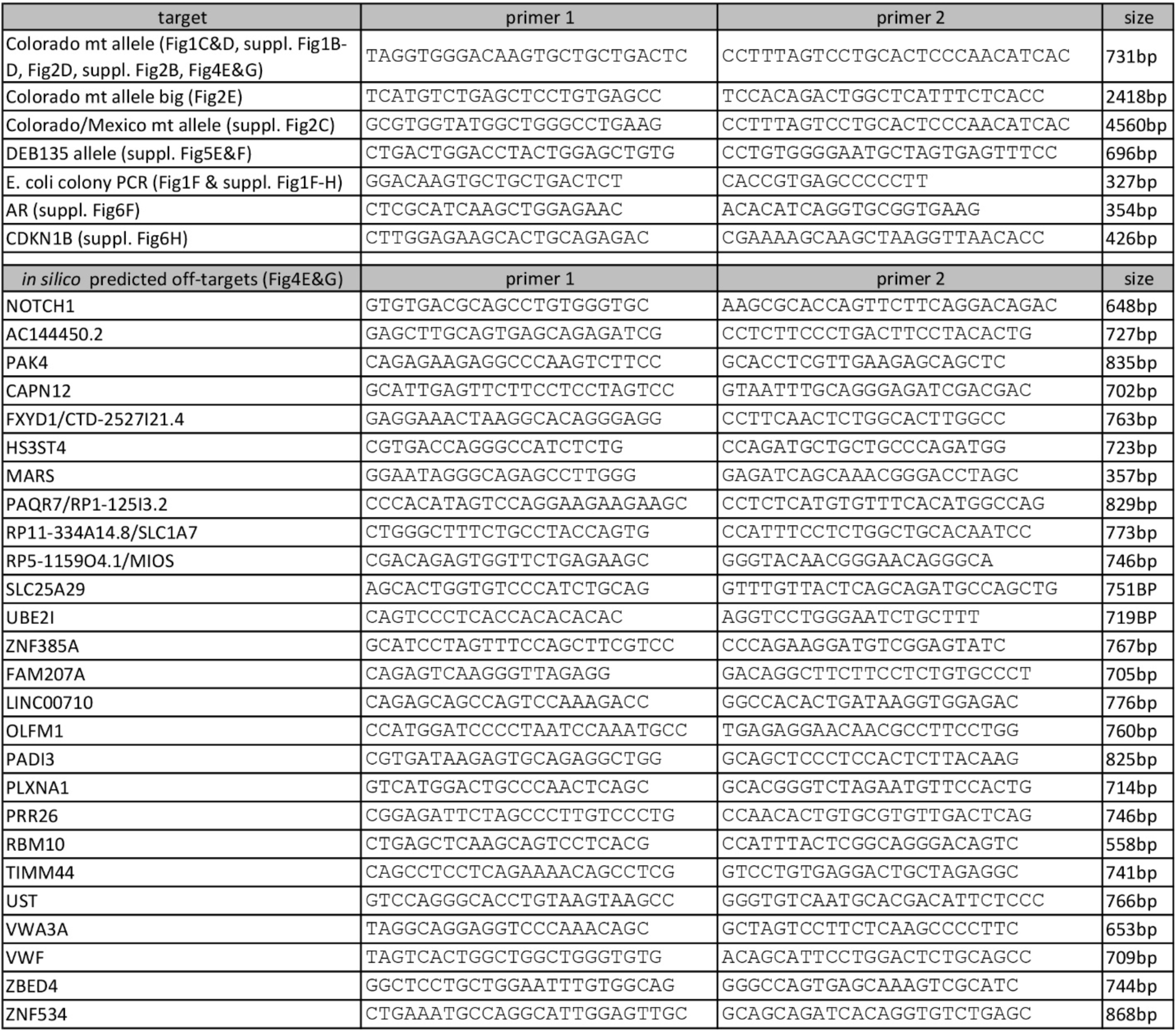
PCR primer sequences.

## References

1. Magrin E, Miccio A, Cavazzana M. Lentiviral and genome-editing strategies for the treatment of beta-hemoglobinopathies. Blood 2019;134(15):1203–1213. DOI: 10.1182/blood.2019000949.

2. Tucci F, Galimberti S, Naldini L, Valsecchi MG, Aiuti A. A systematic review and meta-analysis of gene therapy with hematopoietic stem and progenitor cells for monogenic disorders. Nat Commun 2022;13(1):1315. DOI: 10.1038/s41467-022-28762-2.

3. Lee A, Xu J, Wen Z, Jin P. Across Dimensions: Developing 2D and 3D Human iPSC-Based Models of Fragile X Syndrome. Cells 2022;11(11). DOI: 10.3390/cells11111725.

4. Thomas D, Choi S, Alamana C, Parker KK, Wu JC. Cellular and Engineered Organoids for Cardiovascular Models. Circ Res 2022;130(12):1780–1802. DOI: 10.1161/CIRCRESAHA.122.320305.

5. Trush O, Takasato M. Kidney organoid research: current status and applications. Curr Opin Genet Dev 2022;75:101944. DOI: 10.1016/j.gde.2022.101944.

6. Rao I, Crisafulli L, Paulis M, Ficara F. Hematopoietic Cells from Pluripotent Stem Cells: Hope and Promise for the Treatment of Inherited Blood Disorders. Cells 2022;11(3). DOI: 10.3390/cells11030557.

7. Selvaraj S, Kyba M, Perlingeiro RCR. Pluripotent Stem Cell-Based Therapeutics for Muscular Dystrophies. Trends Mol Med 2019;25(9):803–816. DOI: 10.1016/j.molmed.2019.07.004.

8. Sebastiano V, Zhen HH, Haddad B, et al. Human COL7A1-corrected induced pluripotent stem cells for the treatment of recessive dystrophic epidermolysis bullosa. Sci Transl Med 2014;6(264):264ra163. DOI: 10.1126/scitranslmed.3009540.

9. Doudna JA. The promise and challenge of therapeutic genome editing. Nature 2020;578(7794):229–236. DOI: 10.1038/s41586-020-1978-5.

10. Yamanaka S. Pluripotent Stem Cell-Based Cell Therapy-Promise and Challenges. Cell Stem Cell 2020;27(4):523–531. DOI: 10.1016/j.stem.2020.09.014.

11. Fulchand S, Harris N, Li S, et al. Patient-reported outcomes and quality of life in dominant dystrophic epidermolysis bullosa: A global cross-sectional survey. Pediatr Dermatol 2021;38(5):1198–1201. DOI: 10.1111/pde.14802.

12. Phillips GS, Huang A, Augsburger BD, et al. A retrospective analysis of diagnostic testing in a large North American cohort of patients with epidermolysis bullosa. J Am Acad Dermatol 2022;86(5):1063–1071. DOI: 10.1016/j.jaad.2021.09.065.

13. Tang JY, Marinkovich MP, Lucas E, et al. A systematic literature review of the disease burden in patients with recessive dystrophic epidermolysis bullosa. Orphanet J Rare Dis 2021;16(1):175. DOI: 10.1186/s13023-021-01811-7.

14. Bardhan A, Bruckner-Tuderman L, Chapple ILC, et al. Epidermolysis bullosa. Nat Rev Dis Primers 2020;6(1):78. DOI: 10.1038/s41572-020-0210-0.

15. Has C, Bauer JW, Bodemer C, et al. Consensus reclassification of inherited epidermolysis bullosa and other disorders with skin fragility. Br J Dermatol 2020;183(4):614–627. DOI: 10.1111/bjd.18921.

16. Shinkuma S, Guo Z, Christiano AM. Site-specific genome editing for correction of induced pluripotent stem cells derived from dominant dystrophic epidermolysis bullosa. Proc Natl Acad Sci U S A 2016;113(20):5676–81. DOI: 10.1073/pnas.1512028113.

17. Hirsch T, Rothoeft T, Teig N, et al. Regeneration of the entire human epidermis using transgenic stem cells. Nature 2017;551(7680):327–332. DOI: 10.1038/nature24487.

18. Kocher T, Petkovic I, Bischof J, Koller U. Current developments in gene therapy for epidermolysis bullosa. Expert Opin Biol Ther 2022:1–14. DOI: 10.1080/14712598.2022.2049229.

19. Siprashvili Z, Nguyen NT, Gorell ES, et al. Safety and Wound Outcomes Following Genetically Corrected Autologous Epidermal Grafts in Patients With Recessive Dystrophic Epidermolysis Bullosa. JAMA 2016;316(17):1808–1817. DOI: 10.1001/jama.2016.15588.

20. Mavilio F, Pellegrini G, Ferrari S, et al. Correction of junctional epidermolysis bullosa by transplantation of genetically modified epidermal stem cells. Nat Med 2006;12(12):1397–402. DOI: 10.1038/nm1504.

21. Rashidghamat E, McGrath JA. Novel and emerging therapies in the treatment of recessive dystrophic epidermolysis bullosa. Intractable Rare Dis Res 2017;6(1):6–20. DOI: 10.5582/irdr.2017.01005.

22. Eichstadt S, Barriga M, Ponakala A, et al. Phase 1/2a clinical trial of gene-corrected autologous cell therapy for recessive dystrophic epidermolysis bullosa. JCI Insight 2019;4(19). DOI: 10.1172/jci.insight.130554.

23. Blanco E, Izotova N, Booth C, Thrasher AJ. Immune Reconstitution After Gene Therapy Approaches in Patients With X-Linked Severe Combined Immunodeficiency Disease. Front Immunol 2020;11:608653. DOI: 10.3389/fimmu.2020.608653.

24. Uchiyama T, Takahashi S, Nakabayashi K, et al. Nonconditioned ADA-SCID gene therapy reveals ADA requirement in the hematopoietic system and clonal dominance of vector-marked clones. Mol Ther Methods Clin Dev 2021;23:424–433. DOI: 10.1016/j.omtm.2021.10.003.

25. Jackow J, Guo Z, Hansen C, et al. CRISPR/Cas9-based targeted genome editing for correction of recessive dystrophic epidermolysis bullosa using iPS cells. Proc Natl Acad Sci U S A 2019. DOI: 10.1073/pnas.1907081116.

26. Kogut I, McCarthy SM, Pavlova M, et al. High-efficiency RNA-based reprogramming of human primary fibroblasts. Nat Commun 2018;9(1):745. DOI: 10.1038/s41467-018-03190-3.

27. Gunschmann C, Stachelscheid H, Akyuz MD, et al. Insulin/IGF-1 controls epidermal morphogenesis via regulation of FoxO-mediated p63 inhibition. Dev Cell 2013;26(2):176–87. DOI: 10.1016/j.devcel.2013.05.017.

28. Kelly OG, Melton DA. Induction and patterning of the vertebrate nervous system. Trends Genet 1995;11(7):273–8. DOI: 10.1016/s0168-9525(00)89074-5.

29. Li A, Lai YC, Figueroa S, et al. Deciphering principles of morphogenesis from temporal and spatial patterns on the integument. Dev Dyn 2015;244(8):905–20. DOI: 10.1002/dvdy.24281.

30. Yamaguchi Y, Itami S, Watabe H, et al. Mesenchymal-epithelial interactions in the skin: increased expression of dickkopf1 by palmoplantar fibroblasts inhibits melanocyte growth and differentiation. J Cell Biol 2004;165(2):275–85. DOI: 10.1083/jcb.200311122.

31. Collier A, Liu A, Torkelson J, et al. Gibbin mesodermal regulation patterns epithelial development. Nature 2022;606(7912):188–196. DOI: 10.1038/s41586-022-04727-9.

32. Lee J, Rabbani CC, Gao H, et al. Hair-bearing human skin generated entirely from pluripotent stem cells. Nature 2020;582(7812):399–404. DOI: 10.1038/s41586-020-2352-3.

33. Tadeu AM, Horsley V. Notch signaling represses p63 expression in the developing surface ectoderm. Development 2013;140(18):3777–86. DOI: 10.1242/dev.093948.

34. Hayashi R, Yamato M, Saito T, et al. Enrichment of corneal epithelial stem/progenitor cells using cell surface markers, integrin alpha6 and CD71. Biochem Biophys Res Commun 2008;367(2):256–63. DOI: 10.1016/j.bbrc.2007.12.077.

35. Metral E, Bechetoille N, Demarne F, Rachidi W, Damour O. alpha6 Integrin (alpha6(high))/Transferrin Receptor (CD71)(low) Keratinocyte Stem Cells Are More Potent for Generating Reconstructed Skin Epidermis Than Rapid Adherent Cells. Int J Mol Sci 2017;18(2). DOI: 10.3390/ijms18020282.

36. Gupta K, Levinsohn J, Linderman G, et al. Single-Cell Analysis Reveals a Hair Follicle Dermal Niche Molecular Differentiation Trajectory that Begins Prior to Morphogenesis. Dev Cell 2019;48(1):17–31 e6. DOI: 10.1016/j.devcel.2018.11.032.

37. Ji AL, Rubin AJ, Thrane K, et al. Multimodal Analysis of Composition and Spatial Architecture in Human Squamous Cell Carcinoma. Cell 2020;182(2):497–514 e22. DOI: 10.1016/j.cell.2020.05.039.

38. Kawakami T, Okano T, Takeuchi S, et al. Approach for the Derivation of Melanocytes from Induced Pluripotent Stem Cells. J Invest Dermatol 2018;138(1):150–158. DOI: 10.1016/j.jid.2017.07.849.

39. Lin Z, Jin S, Chen J, et al. Murine interfollicular epidermal differentiation is gradualistic with GRHL3 controlling progression from stem to transition cell states. Nat Commun 2020;11(1):5434. DOI: 10.1038/s41467-020-19234-6.

40. Wang S, Drummond ML, Guerrero-Juarez CF, et al. Single cell transcriptomics of human epidermis identifies basal stem cell transition states. Nat Commun 2020;11(1):4239. DOI: 10.1038/s41467-020-18075-7.

41. Barrandon Y, Green H. Three clonal types of keratinocyte with different capacities for multiplication. Proc Natl Acad Sci U S A 1987;84(8):2302–6. DOI: 10.1073/pnas.84.8.2302.

42. Enzo E, Secone Seconetti A, Forcato M, et al. Single-keratinocyte transcriptomic analyses identify different clonal types and proliferative potential mediated by FOXM1 in human epidermal stem cells. Nat Commun 2021;12(1):2505. DOI: 10.1038/s41467-021-22779-9.

43. Gupta N, Susa K, Yoda Y, Bonventre JV, Valerius MT, Morizane R. CRISPR/Cas9- based Targeted Genome Editing for the Development of Monogenic Diseases Models with Human Pluripotent Stem Cells. Curr Protoc Stem Cell Biol 2018;45(1):e50. DOI: 10.1002/cpsc.50.

44. Aprilyanto V, Aditama R, Tanjung ZA, Utomo C, Liwang T. CROP: a CRISPR/Cas9 guide selection program based on mapping guide variants. Sci Rep 2021;11(1):1504. DOI: 10.1038/s41598-021-81297-2.

45. Akcakaya P, Bobbin ML, Guo JA, et al. In vivo CRISPR editing with no detectable genome-wide off-target mutations. Nature 2018;561(7723):416–419. DOI: 10.1038/s41586-018-0500-9.

46. Tsai SQ, Nguyen NT, Malagon-Lopez J, Topkar VV, Aryee MJ, Joung JK. CIRCLE-seq: a highly sensitive in vitro screen for genome-wide CRISPR-Cas9 nuclease off-targets. Nat Methods 2017;14(6):607–614. DOI: 10.1038/nmeth.4278.

47. Cai ED, Kim J. Identification of novel targetable mutations in metastatic anorectal melanoma by next-generation sequencing. JAAD Case Rep 2017;3(6):539–541. DOI: 10.1016/j.jdcr.2017.09.036.

48. Callahan CA, Ofstad T, Horng L, et al. MIM/BEG4, a Sonic hedgehog-responsive gene that potentiates Gli-dependent transcription. Genes Dev 2004;18(22):2724–9. DOI: 10.1101/gad.1221804.

49. Melo SP, Lisowski L, Bashkirova E, et al. Somatic correction of junctional epidermolysis bullosa by a highly recombinogenic AAV variant. Mol Ther 2014;22(4):725–33. DOI: 10.1038/mt.2013.290.

50. Peng S, Maihle NJ, Huang Y. Pluripotency factors Lin28 and Oct4 identify a sub- population of stem cell-like cells in ovarian cancer. Oncogene 2010;29(14):2153–9. DOI: 10.1038/onc.2009.500.

51. Jackow J, Rami A, Hayashi R, et al. Targeting the Jak/Signal Transducer and Activator of Transcription 3 Pathway with Ruxolitinib in a Mouse Model of Recessive Dystrophic Epidermolysis Bullosa-Squamous Cell Carcinoma. J Invest Dermatol 2021;141(4):942–946. DOI: 10.1016/j.jid.2020.08.022.

52. Matheus F, Raveh T, Oro AE, Wernig M, Drukker M. Is hypoimmunogenic stem cell therapy safe in times of pandemics? Stem Cell Reports 2022;17(4):711–714. DOI: 10.1016/j.stemcr.2022.02.014.

53. Branzei D, Foiani M. Regulation of DNA repair throughout the cell cycle. Nat Rev Mol Cell Biol 2008;9(4):297–308. DOI: 10.1038/nrm2351.

54. Adikusuma F, Piltz S, Corbett MA, et al. Large deletions induced by Cas9 cleavage. Nature 2018;560(7717):E8–E9. DOI: 10.1038/s41586-018-0380-z.

55. Cullot G, Boutin J, Toutain J, et al. CRISPR-Cas9 genome editing induces megabase-scale chromosomal truncations. Nat Commun 2019;10(1):1136. DOI: 10.1038/s41467-019-09006-2.

56. Kosicki M, Tomberg K, Bradley A. Repair of double-strand breaks induced by CRISPR- Cas9 leads to large deletions and complex rearrangements. Nat Biotechnol 2018;36(8):765–771. DOI: 10.1038/nbt.4192.

57. Liu M, Zhang W, Xin C, et al. Global detection of DNA repair outcomes induced by CRISPR-Cas9. Nucleic Acids Res 2021;49(15):8732–8742. DOI: 10.1093/nar/gkab686.

58. Turchiano G, Andrieux G, Klermund J, et al. Quantitative evaluation of chromosomal rearrangements in gene-edited human stem cells by CAST-Seq. Cell Stem Cell 2021;28(6):1136–1147 e5. DOI: 10.1016/j.stem.2021.02.002.

59. Yin J, Liu M, Liu Y, et al. Optimizing genome editing strategy by primer-extension-mediated sequencing. Cell Discov 2019;5:18. DOI: 10.1038/s41421-019-0088-8.

60. Zuccaro MV, Xu J, Mitchell C, et al. Allele-Specific Chromosome Removal after Cas9 Cleavage in Human Embryos. Cell 2020;183(6):1650–1664 e15. DOI: 10.1016/j.cell.2020.10.025.

61. Li L, Wang Y, Torkelson JL, et al. TFAP2C- and p63-Dependent Networks Sequentially Rearrange Chromatin Landscapes to Drive Human Epidermal Lineage Commitment. Cell Stem Cell 2019;24(2):271–284 e8. DOI: 10.1016/j.stem.2018.12.012.

62. Pattison JM, Melo SP, Piekos SN, et al. Retinoic acid and BMP4 cooperate with p63 to alter chromatin dynamics during surface epithelial commitment. Nat Genet 2018;50(12):1658–1665. DOI: 10.1038/s41588-018-0263-0.

63. Schultz L, Patel S, Davis KL, et al. Identification of dual positive CD19+/CD3+ T cells in a leukapheresis product undergoing CAR transduction: a case report. J Immunother Cancer 2020;8(2). DOI: 10.1136/jitc-2020-001073.

64. Werner B, Case J, Williams MJ, et al. Measuring single cell divisions in human tissues from multi-region sequencing data. Nat Commun 2020;11(1):1035. DOI: 10.1038/s41467-020-14844-6.

65. Brinkman EK, Chen T, Amendola M, van Steensel B. Easy quantitative assessment of genome editing by sequence trace decomposition. Nucleic Acids Res 2014;42(22):e168. DOI: 10.1093/nar/gku936.

66. Concordet JP, Haeussler M. CRISPOR: intuitive guide selection for CRISPR/Cas9 genome editing experiments and screens. Nucleic Acids Res 2018;46(W1):W242–W245. DOI: 10.1093/nar/gky354.

67. Doench JG, Fusi N, Sullender M, et al. Optimized sgRNA design to maximize activity and minimize off-target effects of CRISPR-Cas9. Nat Biotechnol 2016;34(2):184–191. DOI: 10.1038/nbt.3437.

68. 68. Martin M. Cutadapt removes adapter sequences from high-throughput sequencing reads. 2011 2011;17(1):3. (next generation sequencing; small RNA; microRNA; adapter removal). DOI: 10.14806/ej.17.1.200.

69. Li H, Durbin R. Fast and accurate short read alignment with Burrows-Wheeler transform. Bioinformatics 2009;25(14):1754–60. DOI: 10.1093/bioinformatics/btp324.

70. McKenna A, Hanna M, Banks E, et al. The Genome Analysis Toolkit: a MapReduce framework for analyzing next-generation DNA sequencing data. Genome Res 2010;20(9):1297–303. DOI: 10.1101/gr.107524.110.

71. Poplin R, Ruano-Rubio V, DePristo MA, et al. Scaling accurate genetic variant discovery to tens of thousands of samples. bioRxiv 2018:201178. DOI: 10.1101/201178.

72. Cibulskis K, Lawrence MS, Carter SL, et al. Sensitive detection of somatic point mutations in impure and heterogeneous cancer samples. Nat Biotechnol 2013;31(3):213–9. DOI: 10.1038/nbt.2514.

73. Wilm A, Aw PP, Bertrand D, et al. LoFreq: a sequence-quality aware, ultra-sensitive variant caller for uncovering cell-population heterogeneity from high-throughput sequencing datasets. Nucleic Acids Res 2012;40(22):11189–201. DOI: 10.1093/nar/gks918.

74. Fang H, Bergmann EA, Arora K, et al. Indel variant analysis of short-read sequencing data with Scalpel. Nat Protoc 2016;11(12):2529–2548. DOI: 10.1038/nprot.2016.150.

75. Sherry ST, Ward M, Sirotkin K. dbSNP-database for single nucleotide polymorphisms and other classes of minor genetic variation. Genome Res 1999;9(8):677–9. (https://www.ncbi.nlm.nih.gov/pubmed/10447503).

76. Ashburner M, Ball CA, Blake JA, et al. Gene ontology: tool for the unification of biology. The Gene Ontology Consortium. Nat Genet 2000;25(1):25–9. DOI: 10.1038/75556.

77. Gene Ontology C. The Gene Ontology resource: enriching a GOld mine. Nucleic Acids Res 2021;49(D1):D325–D334. DOI: 10.1093/nar/gkaa1113.

